# Top-down engineering of complex communities by directed evolution

**DOI:** 10.1101/2020.07.24.214775

**Authors:** Chang-Yu Chang, Jean C.C. Vila, Madeline Bender, Richard Li, Madeleine C. Mankowski, Molly Bassette, Julia Borden, Stefan Golfier, Paul G. Sanchez, Rachel Waymack, Xinwen Zhu, Juan Diaz-Colunga, Sylvie Estrela, Maria Rebolleda-Gomez, Alvaro Sanchez

## Abstract

Directed evolution has been used for decades to engineer biological systems from the top-down. Generally, it has been applied at or below the organismal level, by iteratively sampling the mutational landscape in a guided search for genetic variants of higher function. Above the organismal level, a small number of studies have attempted to artificially select microbial communities and ecosystems, with uneven and generally modest success. Our theoretical understanding of artificial ecosystem selection is still limited, particularly for large assemblages of asexual organisms, and we know little about designing efficient methods to direct their evolution. To address this issue, we have developed a flexible modeling framework that allows us to systematically probe any arbitrary selection strategy on any arbitrary set of communities and selected functions, in a wide range of ecological conditions. By artificially selecting hundreds of *in-silico* microbial metacommunities under identical conditions, we examine the fundamental limits of the two main breeding methods used so far, and prescribe modifications that significantly increase their power. We identify a range of directed evolution strategies that, particularly when applied in combination, are better suited for the top-down engineering of large, diverse, and stable microbial consortia. Our results emphasize that directed evolution allows an ecological structure-function landscape to be navigated in search for dynamically stable and ecologically and functionally resilient high-functioning communities.

## INTRODUCTION

Harnessing the functions of microbial communities is a major aspiration of modern biology, with implications in fields as diverse as medicine, biotechnology, and agriculture [1]. To accomplish this goal, several groups have successfully engineered small synthetic communities from the bottom-up to carry out functions such as biodegrading environmental contaminants [2–4], manipulating plant phenotypes [5], or producing biofuels [6,7], among many others [8,9]. Despite these and other success stories, the bottom-up engineering of synthetic consortia remains challenging. The function of a consortium is generally affected by species interactions, which are difficult to predict from first principles and grow rapidly with species richness [10–16]. Perhaps more importantly, microbial communities are rapidly evolving ecological systems, and their engineered functions can be disrupted by environmental fluctuations, invasive species, species extinctions, or the fixation of mutant genotypes [17–20].

Rather than fighting these eco-evolutionary forces, an alternative “top-down engineering” approach seeks to leverage ecology and evolution to find microbial consortia with desirable attributes [20–26]. Most of this work has focused on enrichment screens, often followed by dilution or reconstitution to simplify the final community [22,25–28]. Intriguingly, a number of studies have gone a step beyond and empirically demonstrated that ecological communities can respond to artificial selection that is applied at the level of the community itself [29–31]. This strategy has been deployed to iteratively optimize complex microbial communities that modulate plant phenotypes [1,31–35], animal development [36], or the physico-chemical composition of the environment [37–41]. Despite its conceptual elegance, the success of artificial selection at the microbiome level has been mixed and generally modest, and artificial selection has not yet been widely adopted as a means of engineering ecosystem function [1,42].

A limiting factor is that we do not really know yet how to design an efficient artificial selection protocol when the units of selection are microbial communities. The selection methods used in early studies (e.g. [31,43]) were inspired by even earlier work on artificial group selection of either single-species populations [44–46], or two-species communities of sexually reproducing animals [29,47]. In these studies, new generations of communities were created through either: (i) a sexual reproduction-like “migrant-pool” strategy, where the communities with the highest function were mixed together and then used to inoculate a new generation (Fig. 1B), or (ii): an asexual-like “propagule” reproduction strategy, where the best communities were selected and then used as the inoculum for the next generation by propagating them without mixing (Fig. S1) [29,31,37]. All subsequent microbial ecosystem-selection studies followed suit and employed variations of those two methods.

**Figure 1.**
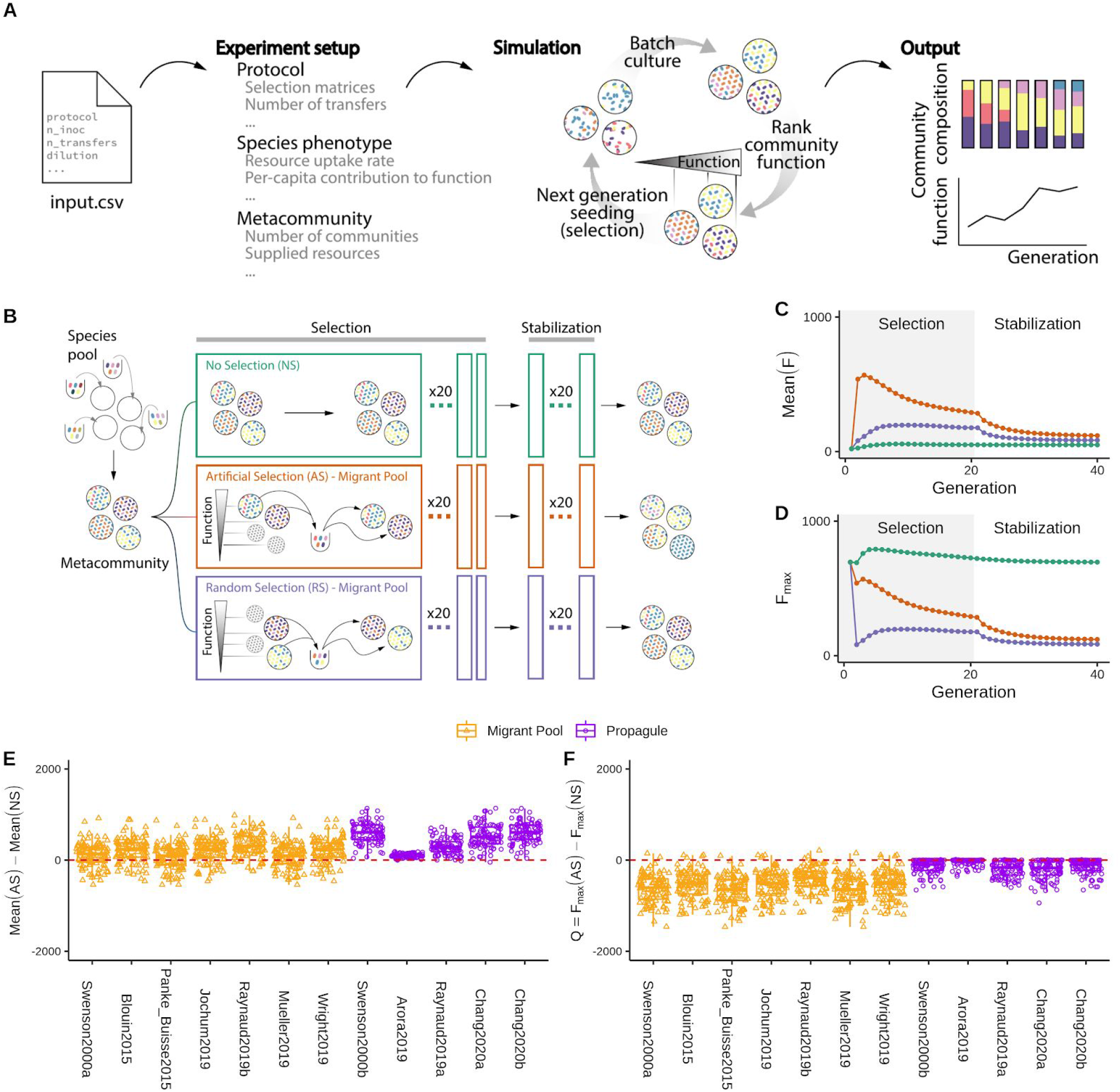
Migrant-pool and propagule strategies are limited in their ability to find new, high-functioning microbial communities. (**A**) We constructed a Python package, *Ecoprospector*, which allows us to artificially select arbitrarily large and diverse *in silico* communities. The experimental design of a selection protocol (e.g., number of communities, growth medium, method of artificial selection, function under selection, etc.) is entered in a single input *.csv* file (Methods). Communities are grown in serial batch-culture, where each transfer into a new habitat is referred to as a community “generation”. Within each batch incubation, species compete for nutrients from the supplied medium. At the end of the incubation period, communities are selected according to the specified, protocol-specific selection scheme, and the selected group is used to seed the communities in the offspring generation. Once the protocol is carried out to completion, *Ecoprospector* outputs a simple text format for later analysis on community function and composition. (**B**) Depiction of a commonly used experimental artificial selection (AS) protocol (the migrant-pool Method [29,44]), alongside its two controls, no selection (NS), and random selection (RS). A representative outcome is shown in (**C-D**), where we adapted the selection protocol from the migrant-pool strategy in ref [31]. A metacommunity is seeded by inoculating 10^6^ randomly drawn cells from a species pool into each of 96 identical habitats and allowing them to grow (Methods). The metacommunity was then subject to 20 rounds of selection (generations), and then allowed to stabilize without selection for another 20 generations. The function maximized under selection *F* is additive on species contributions, whose per-capita species contribution to function is randomly generated (see main text). In each selection round, the top 20% communities with highest *F* (AS; red) (or a randomly chosen set in (RS; blue)) are selected and mixed into a single pool which is then used to seed all communities in the next generation by randomly sampling 10^6^ cells into them. The NS protocol (green) simply propagates the communities in batch mode without selection. The changes in overall function over the generations is shown in (**C**) (average *F*) and (**D**) (maximum function *F*_max_). (**E**) Selection strategies were adapted from twelve experimental protocols in previous studies (see Table S1; Methods). All were applied to standard metacommunity sizes (96 communities), for the same number of generations (20 selection generations + 20 stabilization generations). All protocols have a significantly greater mean function in the AS than in the NS line (Pairwise t-test, P < 0.01) as well as the RS lines (Fig. S4). (**F**) The difference in *F*_max_ between the AS and NS lines (*Q*). All protocols show a Mean *Q* < 0 (Welch’s t-test, P < 0.01), indicating that they did not succeed at improving the function of the best stabilized community in the ancestral population.

But are selection strategies that were originally developed for small populations of sexually reproducing organisms well suited to efficiently direct the evolution of much larger and diverse communities of generally asexual microbes? Are there other alternatives? Here, we investigate a framework for top-down microbial community engineering that is based on the directed exploration of the ecological structure-function landscape (i.e. the map between community composition and community function), through iterated rounds of randomization and selection [10–12,16,48,49]. This approach is inspired by the directed evolution field, where proteins and RNA molecules are evolved in the laboratory through a guided random exploration of their genotype-phenotype maps [50,51]. Unlike molecular fitness landscapes, however, the ecological structure-function landscape is dynamic as species composition changes over time due to inter-species interactions. In what follows, we address how these dynamic landscapes can be navigated, and particularly emphasize the importance of community stabilization before selection.

## RESULTS

### Migrant-pool and propagule breeding strategies are limited in their ability to breed high-functioning microbial communities

To understand the limitations of previous approaches to community-level artificial selection, we first set out to systematically evaluate all previously used variations of the migrant-pool and propagule breeding methods [31,33,34,37–39,52], including two previous experiments from our own group. To do this systematically, one would have to apply all protocols in parallel to the same initial set of communities (hereafter the “metacommunity”[29]), ideally, in multiple replicate experiments and for various different metacommunities. This would require a prohibitively large number of experiments, each with its own control lines. We therefore resorted to *in silico* communities, which can provide the required throughput and allow us to rigorously compare a large number of selection strategies under the same conditions. For that purpose, and inspired by the work of Lenton and Williams [52,53] and others [54–56], we have constructed a flexible computational modeling framework (implemented through a Python package, *Ecoprospector*; Fig. 1A, Methods) that allows us to implement arbitrary community-level selection strategies on arbitrarily large populations of arbitrarily diverse *in silico* communities (Methods). Microbes within a community grow and interact with each other via resource competition following the Microbial Consumer Resource Model (MiCRM) [57–60]. Despite its simplicity, the MiCRM exhibits emergent functional and dynamical behaviors that recapitulate those observed in both natural [59] and experimental communities [61,62].

For simplicity, we have chosen a community function under selection (*F*) that is additive on species contributions: *F*=∑*ϕ*_i_*N*_i_, where *ϕ*_i_ is the per-capita contribution of species *i* and *N*_*i*_ is its population size (Methods). This function is thus redundantly distributed in the community and can be carried out by all single species in isolation. An example of such an additive function could be, for instance, the total biomass of the community, the amount of light scattered on a specific wavelength, or the amount of an enzyme secreted into the environment [10,39,52,54]. Also for simplicity, we assume that the per-capita contribution of a species is fitness neutral. For a recent discussion on the effect of costly functions, see [55]. To start our *in silico* experiments, each metacommunity was seeded by inoculating 96 replicate habitats (all containing the same 90 resources) with N=10^6^ individual cells, which were randomly drawn from 96 “regional pools” of 2,100 species each (Methods). This number of cells is a lower bound for the population sizes that are used in most microbiome experiments (Supplementary Text). Each species is represented by a different randomly sampled vector of nutrient utilization parameters and a randomly sampled per-capita contribution to the function (*ϕ*_i_) (Methods).

Once inoculated, all 96 communities in a metacommunity are allowed to grow for a fixed batch-incubation time *t*, at the end of which we measure their function *F*. A subset of those communities with highest *F* are then selected to breed the next generation, according to the particular selection strategy that is being evaluated (Fig. 1A-B). Strategies were adapted to our specific standardized conditions (e.g. 96 communities, incubation time, dilution factor, etc.) from the papers where they were originally used (Methods; Table S1). To evaluate the effectiveness of these adapted strategies under our *in silico* conditions, we applied each of the twelve selection protocols to the same starting metacommunity for a total of 20 rounds of artificial selection (i.e. community “generations”). To evaluate the stability of the selected function when community-level selection is not constantly applied, we passaged all communities without selection for an additional 20 transfers, giving them time to reach equilibrium (Fig. 1B).

To illustrate a typical outcome, we plot in Fig. 1C-D a representative artificial selection (AS) line where we used the original migrant-pool strategy introduced in ref [31]. For reference, we also show the outcome of a random selection (RS) control, where communities were randomly chosen for reproduction (also adapted from that used in [31]). As shown in Fig. 1C, the mean function in the AS line increases more than in the RS control, indicating a positive response to selection. Importantly, however, the function of the highest-performing community (*F*_max_) in the AS line is lower than in a third “no selection” (NS) control line [44], where each community in the starting metacommunity is simply passaged without community-level selection (Fig. 1D). In other words, a simple “ecological prospecting” procedure, where we screen 96 stable enrichment communities for function and select the best (e.g. [63]) would have found a better community than the multiple rounds of artificial selection we applied at the community level. We note that a NS control line has been missing in all microbiome selection experiments we are aware of.

This experiment illustrates that the mean function in the metacommunity can increase simply because of selection against the worst-performing communities. Importantly, the goal of top-down microbiome engineering is not to improve the mean function in a metacommunity, but to find communities with higher functions than those we started with. Therefore, we propose that the difference between the function *F*_max_ of the highest-performing community (hereafter referred to as the “top community”) in the artificial selection line relative to a no selection control (*Q*=*F*_max_[AS]−*F*_max_[NS]) is a better metric to quantify the success of a selection strategy for top-down engineering purposes. Values of *Q*>0 indicate a successful selection experiment, whereas *Q*<0 indicates an unsuccessful one. Using this metric, we evaluated the success of the twelve propagule and mixed-pool protocols used in previous empirical studies, including our own (Fig. 1E-F) [31–34,36–41]. To obtain a statistically sound assessment, we applied each selection method to N=100 independent artificial selection lines, each with their own NS and RS controls (where applicable; Methods). For all twelve protocols, the mean function increased in response to selection relative to both the NS control (the enrichment screen; Fig. 1E) and the RS control (Fig. S2). Yet, in line with what we observed in Fig. 1D, all protocols failed to improve *F*_max_ (Fig. 1F)

### Selecting communities before they are stable is inefficient

As is the case in all previous artificial selection experiments, our communities are propagated in serial batch-culture. Within each batch incubation, the community goes through an ecological succession. At the end of each batch, a small number of cells are randomly drawn from the community and used to seed a new habitat where all nutrients have been replenished, thus starting a new batch. Inspired by Doulcier et al, we may see each succession as a “developmental” process at the community level, where the communities at the end of a batch incubation can be thought of being in an “adult state” where they are ready for reproduction, whereas the communities at the beginning of a batch incubation are in an “infant state” [54]. In absence of artificial community-level selection, our *in silico* enrichment communities eventually self-assemble into a dynamical state where successions are identical every generation (Fig. S3). Note that this is due exclusively to internal population dynamics, and that no evolution or migration is necessary [61,64–66]. We say that communities are “generationally stable” when the successions are identical across community generations and, therefore, the composition of an adult “offspring” community is the same as that of its adult “parent”. In our simulations, we typically need >5 generations to even approach a generationally stable state.

We speculated that a reason why the selection strategies we evaluated above may be failing to improve *F*_max_ is that, following the original protocols, we started selection at the end of the first batch when the communities are not yet generationally stable. Consistent with this hypothesis, we found that the community rank (from the highest to lowest function) in the first generation is indeed a poor predictor of the function rank of generationally stable communities (Fig. S4A-C; Spearman’s ϱ=0.289, p = 0.004, N=96). In other words, artificial selection in early generations favors communities that had a high function early on, but which end up having mediocre functions once they are generationally stable (Fig. S4). It also selects against those that would have reached a high function in the end, but which started low when they were very far from generational stability. This explains why *Q*<0 for the vast majority of strategies. In the Supplementary Text and Fig. S5-7, we further show that the large population sizes (N=10^6^) of the infant communities in our simulations (which are in the lower end of what is common in experimental microbiome selection) also contribute to the failure of the propagule and mixed pool methods to improve *F*_max_. Both of these methods represent a small subset of all possible ways one could generate an offspring population from the selected members of the parent population. Therefore, we asked if other methods could be used that would increase the success of artificial selection to engineer communities from the top-down.

### An artificial community-level selection strategy inspired by directed evolution of biomolecules

Directed evolution is a form of artificial selection that has been applied for decades to optimize molecular and cellular phenotypes [50,67,68]. In its most common implementation, directed evolution is an iterative process that navigates the genotype-phenotype map in search of a genetic variant of high function [51,69,70]. The process starts by screening a library of genetic variants. Those that are closest to the desired phenotype are selected and their mutational neighborhood is then randomly explored through mutagenesis or recombination, in search of new variants with even higher function. The best among those are selected, and the process can be continued as many times as needed [69]. We reasoned that stable communities of high function can be similarly found through an iterative, guided exploration of their ecological structure-function landscape (Fig. 2A).

**Figure 2.**
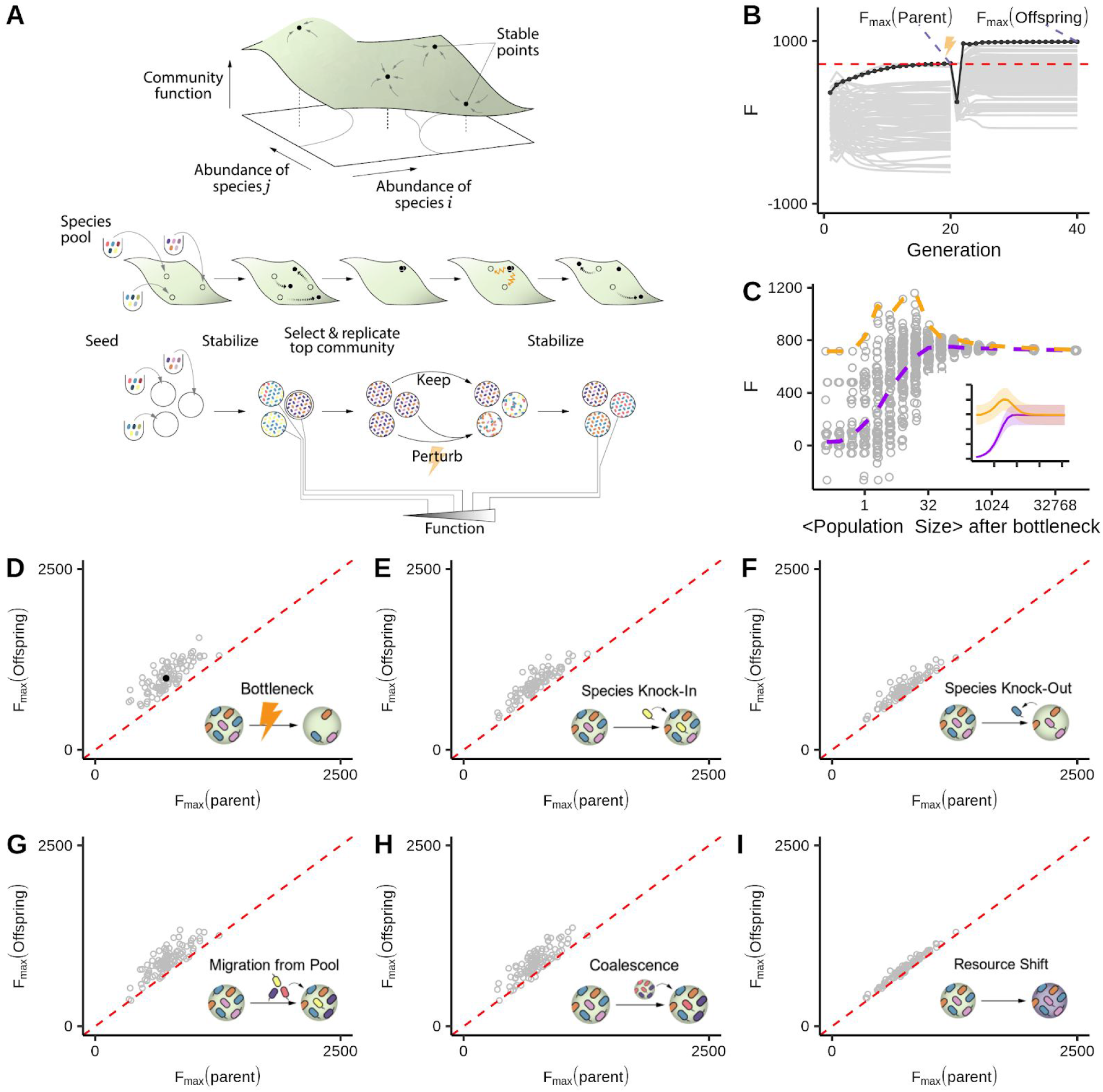
Directed evolution as an artificial selection strategy for high-performing communities. (**A**) Directed evolution of microbial communities can be represented as a guided navigation of a dynamic structure-function landscape, which contains several stable fixed points with different community functions. Community states are given by the abundances of species *i,j* in the “adult” population at the end of a batch. Each of the stable fixed points represents a “generationally stable” equilibrium, as defined in the main text. A library of communities is generated by inoculating from a set of different species pools, followed by stabilization without selection before being scored for function. The top-performing community is selected and either passaged intact into the new generation, or subject to ecological perturbations to generate compositional variants, thereby “exploring” the neighboring stable equilibria in the structure-function landscape. We note that this panel is only a cartoon, the true structure-function landscape is multi-dimensional, and the dynamical stability of equilibria can be significantly more complex than illustrated here. None of those details are critical for our results or discussion. (**B**) A representative outcome of directed evolution of *in silico* microbial communities. N=96 communities are first stabilized by serially passaging without selection with a dilution factor of 10^3^× for 20 generations. The community with the highest function *F*_max_(parent) (black dots and line) is selected, and used to seed the new generation. To that end, the selected community is either passaged intact with the same dilution factor of 10^3^× (N=1), or subject to 95 dilution shocks (10^8^×) to generate variants. The 96 offspring communities are then propagated for another 20 generations until they stabilize. The top offspring variant *F*_max_(offspring) is highlighted with black dots and line. The red dashed line denotes the *F*_max_ of a no-selection line. (**C**) Sampling the optimal bottleneck size by subjecting a single parent community to bottlenecks of different intensity. Each bottleneck is applied 95 times. In orange, we trace the *F*_max_ for the highest-function variants for each bottleneck size. In purple, we track the mean function. Inset shows the outcome of repeating this experiment 100 times with different starting communities (Mean±SD). This shows that intermediate bottlenecks maximize the *F*_max_. (**D-I**) *F*_max_ of 95 stable offspring variants generated through a variety of methods (see text for details), as a function of the *F*_max_ of the (stable) parental community from which all variants were generated. Points above the red dashed line indicated an increase in *F*_max_ from parent to offspring. The filled black circle in panel **D** marks the representative example shown in panel **B**.

A key difference between genotype-phenotype maps and ecological structure-function landscapes is that, whereas the genotype of a gene or even an organism is stable, not all structures (compositions) of an adult community are dynamically stable, and most will either change over time due to drift, or converge to their nearest dynamical attractor (i.e., a generationally stable equilibrium) (Fig. 2A). As we saw in the previous section, it is essential that artificial selection is applied to stable adult communities. Taking into account the dynamical nature of microbial communities, we designed a directed evolution approach which consists of the following sequence of steps (Fig. 2A): (i) An initial library of communities is created by inoculating identical habitats from different species pools, and serially passaged in the absence of (community-level) artificial selection to allow all communities to stabilize. (ii) The stable communities are then screened for a community-level function of interest, and the highest-performing stable community is selected. (iii) The selected stable community continues to be passaged intact into the offspring generation, retaining its function and composition. The rest of the offspring generation will consist of proximal “compositional variants” of the selected parental community. (iv) The offspring communities are allowed to reach their own stable equilibria; and (v) the now stable offspring communities are scored for function (Fig. 2A). The process can be repeated as many times as needed.

How may we generate proximal compositional variants of the best parental community? In Fig. 1, the large population sizes of infant communities (N~10^6^) led to a low between-population variation in the selected function (Supplementary Text), thus failing to generate a diverse enough pool of proximal compositional variants (Fig. S5). We reasoned that with a more stringent propagule bottleneck we may be able to better explore the structure-function landscape, a process that has been successfully applied in the past to converge on simpler, functional consortia by dilution-to-extinction [71]. To test this idea, the top community from a stable parent metacommunity was selected after 20 serial transfers (with dilution factor = 10^3^×), and then used to seed a new generation by subjecting it to 95 separate harsh dilution shocks (dilution factor = 10^8^×), which led to a mean bottlenecked “infant” population size of N=9.76±3.12 cells (Mean±SD) (Fig. 2B). The 95 resulting “offspring” communities differed from each other in which species from the parental community were randomly sampled in the dilution shock. When subject to serial batch culture, they converged to different generationally stable compositions after 20 additional generations (Fig S8). Since they vary in their composition (they are compositional variants), the stable offspring communities also had different functions and some were higher than their parent’s (Fig. 2B).

Consistent with our hypothesis, (Fig. 2C), the propagule method works best at exploring the structure-function landscape and improving *F*_max_ when we use harsh bottlenecks (starting population sizes of order N~10) but it fails at exploring the landscape and generate sufficient between-population variance (therefore not finding a variant with higher *F*_max_) when the number of cells after the bottleneck is above N~10^3^. For mean bottleneck sizes around N~1, community diversity is too low and the function diminishes. The hump-shaped dependence of *F*_max_ on the dilution strength shown in Fig. 2C is consistent with the findings of dilution-to-stimulation experiments [71].

Besides bottlenecks, many other community perturbations can be applied to explore the proximal ecological space in search for compositional variants with high function. For instance, we could create these variants by invading the top parental community with a single, high-*ϕ*_i_ species (i.e. a “knock-in”) (Fig. 2E) [72]. One could also create variants of a community by selectively eliminating (“knocking-out”) one of its members (e.g. a species with low-*ϕ*_i_, or one which competes with a higher-contributing species). In practice, this could be accomplished using narrow-spectrum antibiotics, bacteriophages, or other targeted elimination approaches (Fig. 2F) [73–75]. Larger and more blunt perturbations are also possible: for instance, we could create a library of variants by invading the top parental community with a randomly sampled set of species from multiple different regional pools (i.e. a set of migration events) (Fig. 2G), or by coalescing the top parental community with a library of other stable communities [76,77] (which is a form of migrant-pool method) (Fig. 2H). Another approach is to introduce a library of small random shifts to the nutrient composition, which leads to a rearrangement in species composition and therefore to different function values (Fig. 2I). We have applied all of these perturbations to N=100 independent lines (Methods), and in all cases they were successful at producing one or more dynamically stable community variants with higher function than the best member of the parent population (Figs 2D-I).

### Iteratively combining bottlenecks and migrations to optimize community function selects for high-functioning communities that are ecologically stable

Some of the perturbations in Fig. 2 work by eliminating taxa that are deleterious to community function (e.g. the single species knockouts or the dilution shocks). Others work by adding taxa that are beneficial to community function (e.g. single species knock-ins, or multi-species invasion from the regional pool). We hypothesized that a method that combines random elimination of resident strains with random addition of new strains could help us find high-function variants, as such a method could simultaneously eliminate deleterious species and add beneficial ones. Although random deletion can also eliminate beneficial strains and random addition may contribute deleterious ones, by “tossing the dice” a sufficiently large number of times we have a chance to find one or more variants where the combined effect of species eliminations and additions aligns in the same beneficial direction, reaching high-function regions of the ecological space that we could not find through either method alone.

To test this hypothesis, we directed the evolution of a metacommunity (N=96 communities) using either a species deletion protocol (the dilution bottleneck in Fig. 2D), a species addition protocol (the migration we introduced in Fig. 2G), or a protocol that combined both perturbations simultaneously (Fig. 3A; Methods). As we show in Fig. 3B-D, the strategy that combines both perturbations finds a higher-function community than any of the two perturbations alone. When we replicated this experiment 100 times with as many different metacommunities, we found that the combination of both perturbations produced a significantly higher *Q* than either the dilution shock (*Q* = 641±162 vs 155±94, Mean±SD, paired t-test, p<0.01, N=100) or the migration protocol (*Q* = 641±162 vs 437±152, Mean±SD, paired t-test, p<0.01, N=100) alone (Fig. 3E).

**Figure 3.**
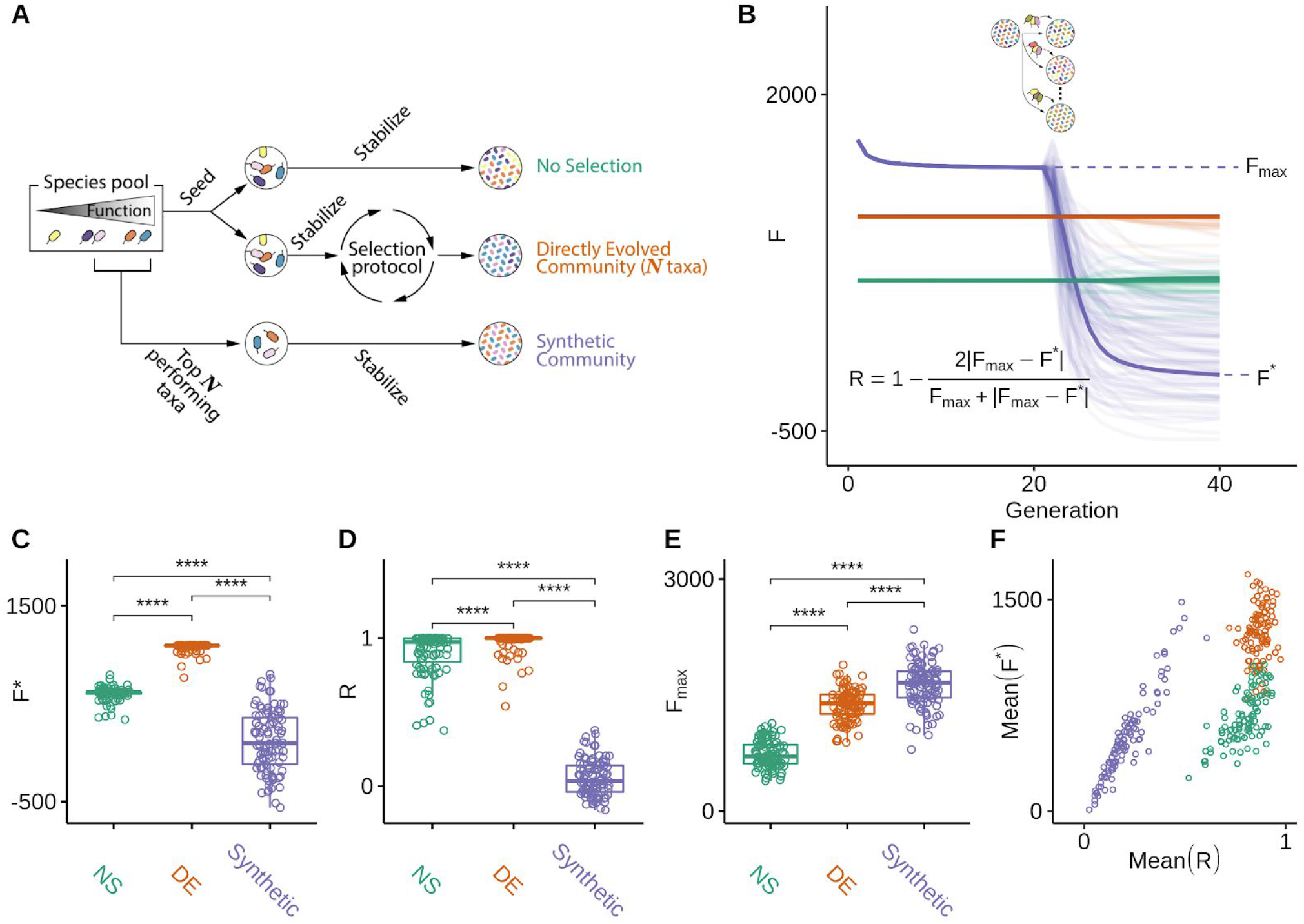
Iteratively combining bottlenecks and migrations to optimize community function selects for high-functioning communities. (**A**) Schematic of iterative protocols of directed evolution. A metacommunity of 96 communities was stabilized for 30 generations by serial batch-culture with dilution factor 10^3^×(Methods). The top community after 30 generations was selected, and either passaged intact to the offspring generation, or used to generate 95 new variants by three different means: (red) in addition to the regular dilution factor (10^3^×), we applied a harsh bottleneck (10^4^×); (purple) we applied a migration event where 10^2^ cells (~45 species; Methods) were randomly sampled from the regional pool and added to each community immediately after passaging them with the regular dilution factor of 10^3^×; (green) a combination of both: after the passage with regular dilution factor (10^3^×), communities are first bottlenecked with a dilution factor (10^4^×), followed by migration from the regional pool (10^2^ cells of ~45 species). The 96 offspring communities are stabilized for an additional 20 transfers, following which they are scored for function. The process can be iterated at this point (**B-D**) *F* for all communities in each generation as a function of time. Each vertical dashed line marks the time points at which the metacommunities experience selection followed by generation of new variants (color represents perturbation type). Red horizontal lines represent the *F*_max_ of a no-selection line. (**E**) *Q* obtained from each of the three protocols at the final time point (generation 460) in n=100 independent selection lines Each point represents the outcome of a different directed evolution experiment. Brackets represent paired t-tests (N=100 for each test). ****:p<0.0001.

An important strength of using directed evolution to engineer microbial communities from the top-down is that we find high-functioning communities that are dynamically stable. Because, by design, the function we are selecting for is additive and the per-capita contribution of each species (*ϕ*_i_) is not affected by other species, one could argue that a “synthetic” bottom-up approach where we just mix together high-contributing taxa would have likely worked just as well (if not better) than our artificial selection protocols. While this may be true, we also reasoned that since the communities in Fig. 3 have been formed by recurrent invasions from the regional species pool, they are also likely to be more robust to external invasions than “synthetic” bottom-up consortia. To test this hypothesis, we went back to the artificial selection line shown in Fig. 3D, and created a “bottom-up” synthetic consortium by mixing together the *n*-th species with the highest *ϕ*_i_ in the regional pool (where *n=*12 is the number of taxa in our artificially selected community, allowing us to control for the effect of biodiversity on functional stability [78]) (Fig. 4A; Methods). We then allowed this synthetic community to stabilize in the same environment and propagation conditions that were used in the artificial selection line.

**Figure 4.**
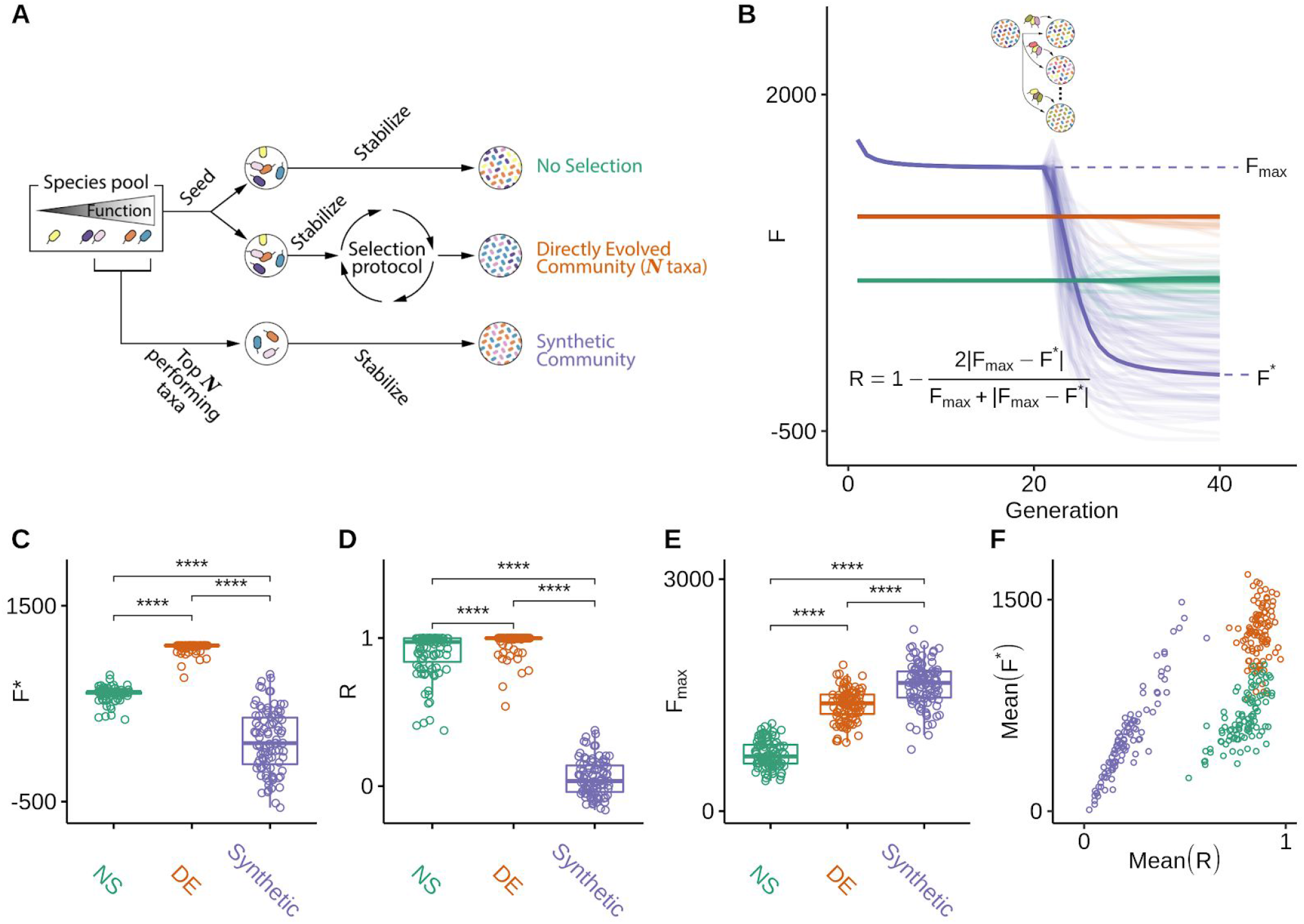
Directed evolution produces communities that are resistant to ecological perturbations. **(A)** We compare the function and ecological stability of communities engineered from the top-down by directed evolution (DE; red), with a synthetic bottom-up consortium (purple). A no-selection (NS; green) control is also provided for reference. The DE community was found by multiple rounds of selection using a protocol that combines bottleneck and migration to generate variants. The synthetic community of equal diversity (species richness) was assembled by mixing together high-ɸ_i_ species from the regional pool (Methods). The top community of the NS control was also chosen as reference. (**B**) The three communities were stabilized for 20 generations (note that the DE and NS were already in equilibrium at the start, but the synthetic community was not). After that, each community was subject to invasion by a randomly sampled set of species from the regional species pool (Methods). This process was repeated 95 independent times for each community. The perturbed communities (lighter-color lines) were allowed to equilibrate by passaging for an additional 20 generations without artificial selection. Following the perturbation, communities reached a new state with function *F*\*, and from the changes in function before and after the perturbation we compute the resistance *R* (inset equation) [79]. (**C-D**) The values of *F*\* and *R* resulting from panel (**B**) are plotted. Values above brackets represent p-values of paired t-tests (N = 95 each test). (**E-F**) The experiment in (B) was repeated 100 times with as many different initial DE, NS, and synthetic consortia. (**E**) shows *F*_max_ of 100 independent experiments. Values above brackets represent p-values of paired t-tests (N=100 each test) (**F**) Mean(*R*) vs Mean(*F*\*) for all 100 independent experiments. *:p<0.05, ****:p<0.0001.

As shown in Fig. 4B, the stable synthetic community has indeed higher function than the directed evolution experiment. Yet, when we invaded both communities with the same random sample of species from the regional pool (containing 100 cells and, on average, 45 species; Methods), the function of the synthetic community collapsed below the artificially selected community (Fig. 4B). By averaging over 95 independent invasion experiments, we obtained the average Resistance (*R*) (a metric of ecological stability, which we calculated as in Shade et al [79]), as well as the average community function after invasion (*F*\*). As we anticipated, the artificially selected community was more resistant to invasion than the synthetic consortium (*F\**=1077±48 vs 114±314, and *R*=0.974±0.05 vs 0.054±0.122, respectively; Mean±SE, p<0.01 in both cases; paired t-test, N=95) (Fig. 4C-D). The synthetic consortium also has lower resistance than the one found by an enrichment screen (the top community in the NS line: *R*=0.896±0.150 vs 0.054±0.122, respectively; Mean±SE, p<0.01; paired t-test, N=95) (Fig. 4C-D).

When we repeated every step of this experiment for the remaining 99 artificial selection lines in Fig. 3E, we found that these results were generic. The function of the bottom-up synthetic consortia (*F*_syn_) is generally higher than the *F*_max_ found through directed evolution and enrichment screens (Fig. 4E). However, the synthetic communities are less resistant to invasion than the artificially selected communities (Mean(*R*)=0.867±0.045 vs 0.214±0.115, and Mean(*F\**)=1261±189 vs 525±307 respectively; Mean±SE, p<0.01 in both cases, paired t-test, N=100) (Fig. 4F). Importantly, the directed evolution communities were more resistant to invasion, on average, than those found through enrichment screens, even though resistance to invasion was not directly selected for (*R* = 0.867±0.045 vs 0.793±0.087 and Mean(*F*\*) =1261±190 vs 660±180; Mean±SE, p<0.01 in both cases, paired t-test, N=100) (Fig. 4F). This indicates that the repeated migrations that are part of the protocol in the directed evolution experiment confer the selected communities with higher stability to this perturbation (though not necessarily to other perturbations, Fig. S9). Our results also suggest that a simple screen may allow us to find a more ecologically stable (if also lower functioning) community than a synthetic consortium, at least when ecological stability is not engineered as well into the consortium.

## DISCUSSION

A growing number of techniques are making it possible to edit the genomes of microbes within microbial communities [74,80,81]. Simple ecological methods to modify microbiome composition also exist, from dilution-to-extinction [22,27], to species engraftment [72]. Together, these tools are paving the way to extend directed evolution from its usual sub-organismal domain to the microbiome level. However, working with large, diverse communities of asexual microbes as the unit of (artificial) selection presents unique challenges and opportunities that set it apart from directed evolution at or below the organismal level. For instance, whereas the genotype-phenotype map of an organism is stable over its lifespan, the ecological structure-function landscape is dynamic, as the composition of the community changes from the moment of inoculation to the time of harvesting. Every new genetic variant will have a phenotype, whereas most new community compositional variants will be unstable and will move toward a different stable composition over time. While fitness landscapes can be climbed through mutation and selection, structure-function landscapes must be navigated, requiring an intermediate stabilization step in between randomization and selection.

The population dynamics within a single batch represent an ecological succession. Recently, they have been elegantly conceptualized as a kind of “developmental maturation”, where a community is in an “infant” state at the time of inoculation, and it is an “adult” at the time of harvest and reproduction [54,55]. When death rates are high, it is possible for communities to reach a steady state within a single succession, thus reaching generational equilibrium by the time they are an adult [54]. When death rates are low, as is the case in our simulations (and our enrichment experiments [61]), then multiple serial passages are needed for communities to reach generational equilibrium. This means that adult communities in early generations can still be moving in their ecological structure-function landscape, and should not be subject to selection in that state. At least one artificial selection study found evidence of an unstable succession between transfers in the early selection rounds [38], and this was consistent with recent reports of functional stabilization taking >6 community-generations in sequential enrichment communities [23,28,61,64].

When we examined the effectiveness of the two main methods of artificial group selection that have been used in the past, the migrant-pool and the propagule methods, we found that both underperform when applied to large microbial ecosystems that are not yet generationally stable (Fig. S4D-E). We note, however, that both strategies worked much better when a strong bottleneck is applied and “infant” population sizes are comparable to those used in animal studies (Fig. S10). Based on our simulations, we suggest that (i) ensuring that communities are generationally stable before selection is applied, and (ii) systematically exploring the effect of bottleneck size on between-community variation, will both enhance the effectiveness of both strategies.

Directed evolution can be used to iteratively optimize the function of microbial communities, through sequential rounds of exploration and selection. Previous approaches to engineer communities from the top-down include enrichment (which is often followed by a perturbation such as a bottleneck, to reduce community complexity) [20,22–24,28,82,83], and selective breeding by artificial selection [1,31–34,36–42]. The directed evolution approach we have studied here combines components of both approaches: the iterative search that is inherent of the latter, with the idea of building stable consortia and exploring compositional variants of the former. In addition to inducing evolutionary changes in the resident species, the methods to generate compositional variants and explore the ecological structure-function landscape include many ecological perturbations that randomly sample new species in and out of the community. For instance, bottlenecking (also known as dilution-to-extinction [21,22,27,71,82,84]) is a blunt method for randomly removing “deleterious” taxa, which has the cost of also eliminating potentially beneficial species. Horizontal immigration from the regional pool may create variants that contain new and potentially “beneficial” species, but it has the cost of potentially adding species with deleterious effects. A selection method that combines the two with strong selection is able to compensate for the specific weaknesses of each, leading us to high-function regions of the ecological structure-function landscape that were not reached by any of the two individual strategies alone (Fig. 4). These communities are also ecologically resistant to invasions compared to both enrichment and synthetic communities assembled by artificially mixing together species with high per-capita function.

Our study has important limitations, as the space of all possible ecological scenarios and methods of artificial community-selection is enormous and we have barely scratched its surface. For instance, we have limited ourselves to rank-based selection, as opposed to assigning communities a reproductive success based on their function, i.e. a “fitness”. Our simulations lack a host, and focus on community-level functions, whereas many of the microbiome selection experiments to date have applied artificial selection based on host traits, rather than direct community-level phenotypes [31–33,36]. Additional work will be needed to extend the *Ecoprospector* package to indirect selection on host traits [33]. In our simulations, species interact exclusively via nutrient competition, excluding the important case of inhibitory and other chemically mediated interactions [85]. The MiCRM can accommodate these interactions organically, and it should be straightforward to implement them in future iterations of this work. For simplicity, we have also assumed that the value of the selected community function does not affect species fitness, and we also focused on a function that is additive on the species contributions and which is not costly at the individual level. We have also not exhaustively considered several practical aspects of artificial community-level selection, which have been studied recently and which are known to be important for the success of this approach [55]. Perhaps the most relevant of these would be the role of non-heritable variation, which may arise from various sources: from pipetting during transfers to day-to-day variations in the environment. These non-heritable sources of between-community variation will all reduce heritability, working against ecosystem-level selection. Along the same lines, our analysis and its discussion has centered in an ecological regime where within-community population dynamics is dominated by selection, with little ecological drift. Since many microbiome selection experiments take place in open ecosystems [31,33,34], it is very likely that one would find species entering and leaving the communities, transiently invading without fixing [86–88]. How these transient species should affect ecosystem-level heritability remains poorly understood. Although the above (and many other) factors have not been considered here, the *Ecoprospector* package we have developed is flexible and we are confident that it can accommodate all of those additional scenarios and many others.

Directed evolution has transformed protein engineering [50], but its application above the organismal level is still in its infancy. Developing methods to create and screen variants was instrumental in the original development of directed evolution in protein engineering [89]. Our stillrudimentary of theoretical understanding of how microbial communities assemble and how they evolve is hampering the design of similarly successful methods to direct the evolution of communities. The modeling framework and computational tools we have introduced in this paper allow us to rapidly test different methods of community selection in a wide range of ecological scenarios. We hope that our results will not only clarify the limitations of previous approaches to artificially selecting communities, but also motivate the development of new methods for the top-down engineering of microbial communities.

## METHODS

### The *Ecoprospector* package

All the results presented in this paper are generated using *Ecoprospector*, a new freely available Python package for implementing artificial selection of microbial communities using customisable protocols. The package builds off the recently published *community-simulator* package (which is a dependency of *Ecoprospector*) [58] and implements protocols in a modular manner that allows an extremely large parameter space of possible protocols to be explored. The parameters for all simulations implemented in this paper are stored in *csv* files (Supplementary Materials).

### Microbial Consumer-Resource Simulations

We model microbial community dynamics using the Microbial Consumer Resource Model (MiCRM) [57–60]. The MiCRM is a minimal model for microbial communities growing in well-mixed resource limited environments (such as in batch culture or in a chemostat). Briefly,the MiCRM models the change through time in i) the abundance of a set of consumer species denoted by *N*_i_ and ii) the concentration of a set of resources denoted by *R*_α_. Ecological interactions between species arise from the uptake and release of resources to the environment. In our simulations consumer and resource dynamics are described by the following sets of equations:

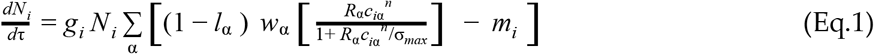

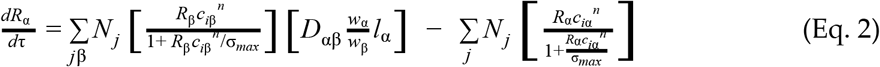

The parameters and units of this model are described and defined in Table S2 (adapted from Marsland et al 2020). This version of the MiCRM assumes that the dependence of resource import rate on resource concentration follows a Hill (type-III) function: 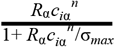. We have also assumed that there is no external resource supply, reflecting resource depletion in a batch-culture like environment. In our simulations, we did not allow cross-feeding *(l=0)* and assumed there is no minimal energy requirement *(m=0)* to eliminate starvation-induced depletions in population size. Aside from the consumer uptake rates (see below) the values of the other MiCRM parameters are set to the default values used in the *community-simulator* package.

### Initial conditions and consumer uptake rates

All simulations are done considering a metacommunity made up of multiple independent communities each of which is simulated using the MiCRM. In order to set the consumer uptake rates across the metacommunity and the initial resource and species compositions of each community, we have adapted the method for constructing random ecosystems from *community-simulator*. The parameters associated with this and the values we used are outlined in Table S3 (adapted and expanded from Marsland et al 2020). Unless otherwise specified these values are set to the default values used in the *community-simulator* package. These parameters will be referenced throughout the rest of the methods section.

### Uptake Rates

Species differ solely in the uptake rate for different resources *c*_*iα*_. For simplicity we assume there is a single family of consumers, a single class of resources and there is no preferential allocation of consumer capacity to different resource classes (*T* = 1, *S*_*f*_ = 1, *q* = 0) (See [57,58] for further explanation of these parameters). Therefore all *c*_*iα*_ are sampled from the same probability distribution. In *community-simulator c*_*iα*_ can be sampled from one of three different distributions: i) a Gaussian distribution ii) a Gamma distribution, or iii) a Bernoulli distribution with binary preference levels set by *c*_0_ and *c*_1_ (referred to as the binary model). Denoting the total uptake capacity of species *i* by 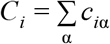, these distributions are parameterized in terms of mean and variance in total uptake rate: μ_*c*_ = < *C*_*i*_ > and σ_*c*_^2^ = *V ar*(*C*_*i*_).

For our purposes none of these distributions were suitable because i) we wanted all *c*_*iα*_ to be positive (unlike the gaussian distribution) ii) we wanted *c*_*iα*_ of some resources to be 0 (unlike the gamma distribution) and iii) we wanted more than two possible values of *c*_*iα*_ (unlike the binary model), To address these limitations, we introduced a new sampling method that combines the gamma distribution and the binary model. Under this approach *c*_*iα*_ can be written as the product of *X* and *Y*, where *X* is sampled using the binary model and *Y* is sampled from a gamma distribution. The mean and variance of *Y* is constrained to values that ensure that mean and variance of *C*_i_ are still μ_*c*_ and σ_*c*_^2^.

Specifically under the binary model :

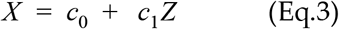

Where *Z* is sampled from the Bernoulli distribution with 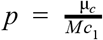, where *M* is the total number of resources (Table S3). Therefore

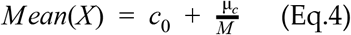

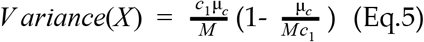

Because *c*_*iα*_ = *XY*, to ensure that μ_*c*_ = < *C*_*i*_ > and σ_*c*_^2^ = *V ar*(*C*_*i*_) we set :

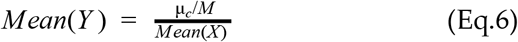

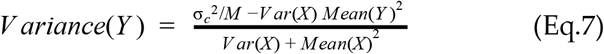

### Initial Resource conditions

All communities within a metacommunity are grown on the same set of *M* = 90 resources. The initial abundance of each resource 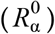 is obtained by first sampling it from a uniform distribution between 0 and 1 and then normalizing 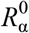 so that the total resource concentration 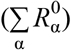 is equal to *R*_*tot*_ = 1000. All communities within a metacommunity have the same 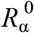 for all α.

### Initial Consumer conditions

Each simulation of a single metacommunity consists of *H* = 2100 species. Each community is inoculated with *n*_*inoc*_ cells sampled (with replacement) from different ‘species pools’ (Table S3). The abundance of each species *i* in the species pool (*U*_*i*_) follows a power-law distribution with exponent *a* = 0.01 and probability density function:

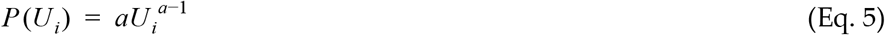

Cell counts are converted into initial species abundances *N*_*i*_ through a conversion factor ψ which we set at 10^6^. This means that 10^6^ cells is equivalent to *N*_*i*_ =1. Because each community is inoculated from a different species pool (each with its own abundance distribution), the abundance of each of the *H* species differs across communities ensuring that our simulations start with compositional and functional variability (Fig 2B).

### Incubation

Once the initial resource and consumer abundance has been set and the MiCRM parameters have been sampled a single community-level ‘generation’ is simulated by propagating the system forward for an incubation time *t* via numerical integration of dynamical equations (Eq. 1–2). During a batch incubation, resources are depleted and consumers increase in density. At the end of each batch incubation (time *t*) the function of each community in the metacommunity is quantified (see following section) and the communities are ranked in terms of their function.

### Community function

The function *F* of each community is measured at the end of each generation In principle any arbitrary community function can be chosen as the “phenotype” under selection. For example one could consider the total biomass of the community, the species richness of the community, the distance of the abiotic environment from some target state, the resistance to invasion etc. In the main text of this paper, we have limited our analysis to a simple additive case:

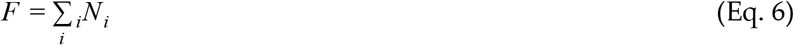

where *ϕ*_i_ is the per-capita contribution of species *i* and *N*_*i*_ is its population size. We sample *ϕ*_i_ from a normal distribution of mean 0 and standard deviation σ = 1.

### Selection matrix

After the function of each community has been measured, the ‘parental’ communities are ‘passaged’ to produce a new set of ‘offspring’ communities. The metacommunity size is kept constant (so the number of offspring communities is equal to the number of parent communities). Passaging simulates the pipetting of bacterial culture into wells containing fresh media (such as from one 96-well plate into another). Which parental communities are selected to contribute species to each offspring community depends on its ranked function. This is specified by a selection matrix *S* whose element *S*_*uv*_ determines the fraction of cells from the parental community of rank function *v* that are used to inoculate offspring community *u* (Fig. S11).

In principle any arbitrary fraction of a parent community of rank *v* can be transferred to offspring community *u*. For the simulations presented in this paper we have set a standard dilution factor *d* and all non-zero elements of the S matrix are set to 1/*d*. Note that the selection matrix *S* is similar to the transfer matrix *f* used in the *community-simulator package [58]* except the indices of the parent community are based on the ranked function of the community rather than being positional.

We also note that whilst for most experiments rank function is determined across the entire metacommunity, for a few simulations we carried out here we used block designs (Fig. S13B). In these cases a metacommunity is divided into multiple sub-metacommunities and the rank function is quantified within each sub-metacommunity. The rank function within each sub-metacommunity is then used to determine which parents are selected. For these cases the selection matrix is divided into blocks along the *v* axis with each sub-metacommunity being allocated one block (Figure S13B). Communities are then sorted by rank along the *v* axis within each block. See [36,40] in Table S1 for examples.

### Passaging

The passaging algorithm considers the transfer of both resources and species. Resource concentrations *R*_α_ are treated as continuous, and we assume they are transferred without any noise. Let *R*^*u*^_α_ be the concentration of resource α in offspring community *u* and *R*^*v*^_α_ be the concentration of resource α in parent community with rank *v*.

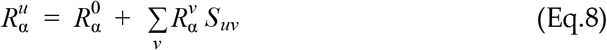

Species abundances *N*_i_ are treated as discrete in order to incorporate demographic noise. Let *N*_i_^*u*^ be the abundance of species *i* in offspring community *u* and *N*_i_^*v*^ be the abundance of species *i* in parent community with rank *v*. For each transfer from community *v* to *u* The total number of cells (of all species) (*z*) passaged is distributed according to a Poisson distribution:

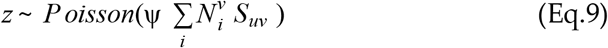

The species identity of each cell transferred to community *u* is then determined by multinomial sampling with the probability of any one cell belonging to species *i* being equal to the relative abundance of species *i* in the parent community 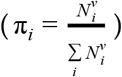. Note that aside from the introduction of Poisson sampling for the total cell number, this procedure is identical to the one used in the *community-simulator package*. Poisson sampling accounts for variability in total number of cells transferred after each generation, an important source of error (compositional variation) in the batch culture lab experiments we are modelling here.

### Random seed

Each metacommunity is associated with a single random seed that is used to determine the initial conditions for the simulation. Each random seed uniquely determines the set of *c*_iα_, *ϕ*_i_, *R*_α_^0^. Whilst each community in the metacommunity will have different initial species abundances, each random seed is associated with a unique set of initial species abundances across the entire metacommunity. As well as ensuring that our results are reproducible, this allows us to carry out different protocols on identical sets of starting communities. For this reason most statistical comparisons in this paper are paired, reflecting the fact that the results are non-independent when different protocols are tested using the same random seed.

### Comparing the effectiveness of all previously used artificial community-level selection protocols using *Ecoprospector*

To systematically evaluate all of the previously used experimental protocols for artificial microbiome selection, we have adapted them into a standardized format that can be simulated using *Ecoprospector*. These protocols, listed in Table S1, were originally designed for microbial communities that had assembled in environments as diverse as the rhizosphere, animal guts, or water treatment plants. Thus, they vary considerably in design. Differences include, the phenotype under selection, the incubation time, dilution factor, number of artificial selection lines, the number of communities per line, and the number of community generations and the controls that were carried out (Table S1).

To standardize the size of all artificial selection lines we consider a metacommunity of 96 communities because: i) this is close to the largest number of communities previously considered in a single artificial selection experiment [41], and ii) 96 well plates are widely used in high-throughput microbial ecology and evolution experiments. To assess these protocols independently of system-specific details, we consider only the selection method used, i.e at the end of each community generation what fraction of communities are selected and how are they transferred, as well as which generation selection starts. This means that our adaptation of each protocol differs solely in the specific 96 × 96 S matrix. We illustrate the S matrices used for all protocols in Table S1.

The microbiome selection experiments we have examined used either a propagule strategy or migrant-pool strategy. For three of five studies that employ propagule strategies (Swenson2000b, Chang 2020a, Chang 2020b) this simply involved selecting a fraction ϱ of communities from the parent generation, and seeding 1/ϱ offspring communities from each of these selected communities.

In the *S* matrix this is encoded by each of the top *ϱ*×96 parent communities being transferred to (different) 1/*ϱ* offspring communities. In cases where *ϱ*×96 is not a whole number, it is rounded up to the nearest integer and all lower function offspring communities are discarded (to ensure metacommunity size remains at 96). For example, if ρ is 0.1, 10 communities will be selected. The 6 top communities will seed 10 offspring communities and the remaining 4 will seed 9 offspring communities. For six of the seven studies that employ migrant-pool strategies this involves selecting a fraction *ϱ* of communities from the parent generation, mixing them together and seeding all communities in the offspring generation from this pool. In the *S* matrix this is encoded by each of the top *ϱ*×96 communities being transferred to every offspring community. In cases where *ϱ*×96 is not a whole number, it is rounded up to the nearest integer. Within each strategy, the various experiment protocols differed in the fraction *ϱ* (ranging from 0.1-0.33).

In two studies, both strategies are carried out with a slight modification in how communities are used to seed the next generation. Specifically Raynaud et al and Arora et al performed artificial community level selection using multiple sub-lines rather than one single line [36,40]. In Raynaud et al, both of the experiments had three sub-lines within an artificial selection line. In the experiment that used a propagule strategy, the top community from each sub-line of the 10 communities is selected and used to seed a sub-line of the next generation. In the experiment using the migrant-pool strategy, the top community from each of the three sub-lines are mixed into a pool that is used to seed all new communities. This is reflected in the selection matrix dividing the 96 communities into three sub-lines of 32 communities each (with only the top member of each sub-line being selected). Arora et al 2019 used a similar propagule strategy, and also used multiple parallel sub-lines this time with each containing three communities. This selection scheme is adapted in our simulation by grouping the communities into 32 sub-lines of three communities.

A number of studies implemented a control strategy involving random selection at each generation [31,32,36,38–41]. For these studies, we also tested the random selection controls by randomizing the rank of all communities, or of communities within a sub-line where applicable. We illustrate one example of a random control S matrix for each protocol in Table S1.

Our simulations are seeded with an initial inoculum size *n*_*inoc*_ = 10^6^ cells. The initial inoculum size was chosen to be comparable with the pioneering work on microbiome selection which includes treatments with two inoculum sizes 0.06g and 6g of soil, which may contain anywhere between 10^6^ − 10^10^ cells [31]. This initial inoculum size gives us communities at the start of our simulations with 225±12 (Mean±SD) species, which is also comparable to previous work (i.e 110-1290 ESVs in Goldford et al 2018). The metacommunity is simulated for 40 community generations with a fixed incubation time (*t=1)* and dilution factor (*d=*10^3^×). This number of generations is equal to the largest number of community generations previously reported [31]. The incubation time and dilution factor are set to ensure that all resources are fully depleted at the end of each incubation and the community reaches stationary phase. The population size during stationary phase as well as the number of generation per incubation are of the same order of magnitude as those in experimental microbial populations (~10 bacterial generations per incubation and a final stationary phase population size of 10^9^). We note that whilst we have kept these parameters constant, they in fact varied substantially in prior community selection studies depending on the empirical systems (e.g., dilution factor ranged from 6× to 250×; incubation time between 48 hours and 35 days; Table S1). One protocol optimized incubation time between batches [38]. While we did not capture this “variable *t*” feature, it is straightforward to do in Ecoprospector.

Unlike previous studies we divide our experiments into two phases each lasting 20 generations. In the first ‘selection’ phase, the protocol-specific selection matrix *S* is applied at the end of each growth cycle. In the second ‘stabilization phase’, communities were passaged without selection, i.e., *S*=(1/*d)I*, where *I* is the identity matrix and *d* the dilution factor, as above. Each protocol can thus be expressed in a sequence of 40 selection matrices which contains 20 consecutive *S* matrices and 20 consecutive (1/*d)I* matrices (Figure S13). The function and composition of all communities at the end of each generation (i.e., during stationary phase) is recorded. We compare the effectiveness of each protocol by applying it (and where applicable its corresponding random selection control) to an identical set of starting communities. We also compare all protocols to a ‘no-selection’ control where all communities are passaged without selection for all generations, i.e *S*=(1/*d)I* for all 40 generations. We repeated all protocols 100 times with 100 different random seeds to obtain a statistically sound sample size.

### Directed evolution

After seeding the metacommunity, we allow all 96 communities to stabilize by propagating them for 20 transfers without selection (using *S*=(1/*d)I*, as we do in the no-selection control). After 20 generations the highest functioning community is selected, and passaged into 96 fresh habitats with a dilution factor of *d*=10^3^× (the same one that had been used during the 20 previous transfers). One of these new communities is left unperturbed. The other 95 copies are all perturbed as described below in an attempt to push the community to a new stable state. After the perturbation, all communities, including the unperturbed community, are grown for 20 generations without selection (*S*=(1/*d)I)* to let them reach a stable state.

We consider six different types of perturbation that are applied to the 95 copies of the top-performing community (Table S5):

- *Bottleneck perturbation.* This approach involves subjecting the 95 communities to an additional dilution step. This is done by repeating the passaging protocol described previously using *S*=(1/*d*_*bot*_)*I* where *d*_*bot*_ is the bottleneck dilution factor. For figures 2B and 2D the *d*_*bot*_=10^5^. An average of N=9.76±3.12 (Mean±SD) cells remain in the community after a bottleneck of this magnitude. In figure 2C we carried out this procedure multiple independent times using a gradient of bottleneck dilution factors (ranging from *d*_*bot*_=10, to *d*_*bot*_=8×10^5^.
- *Species knock-in*. This approach involves introducing a different high-functioning species from the regional pool into each of the 95 communities. A collection of candidate high-performing species is first prepared by growing every single species from the regional pool in monoculture, passaging them in the same batch condition as the communities for 20 serial transfers, and then identifying the top 5% of species (threshold percentile *θ*=0.95) according to their rank function. This gives us a collection of 105 candidate species from the pool of *H* = 2100. We then invaded each of the 95 communities with a different randomly chosen candidate from this set. This is done by introducing 10^3^ cells of the chosen species into each community after they have been diluted. 10^3^ is chosen to minimize the probability of stochastic extinction of the invader.
- *Knock out*. This approach involves eliminating one of the species in the community, so that all offspring communities have *n*-1 species (whereas the parent had *n)*. In each community we delete a different taxa. When *n <95* (as is the case for our simulations) the number of perturbed offspring communities is equal to *n*. The 95-*n* ‘Spare’ communities are left unperturbed.
- *Migration.* This approach involves perturbing the communities by invading with them a random set of species sampled from different regional pools. We use the same approach that we used to initially inoculate the communities. To recap, for each community we added *n*_*mig*_=10^6^ cells randomly sampled from different regional species pools, in which the species abundances are distributed according to Eq. 5. The number of cells introduced via migration is comparable to the number of cells in the communities after the regular batch dilution (~10^6^)
- *Coalescenc*e. This approach involves coalescencing the copies of top-performing communities with the other stable communities in the metacommunity before selection. To do this, the parents metacommunity is not discarded at the point of directed evolution. Instead, it is kept and the offspring metacommunity is grown for a single generation (so that both the parent and offspring metacommunities are in stationary phase). The two metacommunities are then mixed, generating a new metacommunity of coalesced communities. Let *J* be the resource and consumer abundance of the offspring metacommunity and *K* be the resource and consumer abundance of the parent metacommunity. The consumer and resource abundance of the mixed metacommunity *L* is simply:

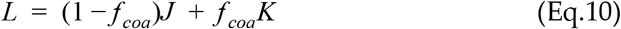 For our simulations we use a *f*_*coa*_ = 0.5 equivalent to mixing equal volumes. To inoculate into the offspring community, the coalesced communities are then diluted with a dilution factor *d*=10^3^× (using *S* = *I/d*).
- *Resource shift*. This approach involves introducing a different random change to the ‘media’ (R_α_^0^) of each of the 95 communities. We have a complex media of M=90 Resources. We first select the most abundant resource in the media and reduce its abundance by δ*R*_1_. This amount of resource is then added to one of the other 89 resources chosen at random. For our simulations we set δ = 1. Unlike other perturbations mentioned above that only happen in the short term (pulse perturbation), the changes in nutrient composition is permanent and persists for the rest of the simulation (press perturbation) [79].

### Iterative directed evolution

To test the effects of iteratively applying directed evolution, we designed a protocol with over 460 community generations that includes 20 consecutive rounds of directed evolution (Figure 3). The protocol starts by seeding the initial metacommunity of 96 communities as before. We grew the communities for 29 generations without selection (*S*=(1/*d*)*I* with *d*=10^3^×). At generation 30 we performed a single round of perturbations (using one or more of the approaches described in the previous section). To reiterate, the top community is selected and passaged (with *d*=10^3^×) into 96 fresh habitats and 95 of these copies are then perturbed to generate compositional variants. The offspring metacommunity is then stabilized for 19 generations without selection after which another round of perturbations is performed. We repeat this sequence of stabilization followed by perturbations 18 more times (giving a total of 20 rounds of directed evolution). After the final round of directed evolution (at generation 410) the metacommunity is grown for 50 generations without selection to ensure it reaches equilibrium.

We simulate this extended protocol using one of three different types of perturbation.

1. Bottlenecks: After the top community has been selected and replicated using a standard dilution factor (*d*) 95 of 96 communities are subject to an additional bottleneck (*d*_*bot*_ =10^4^). An average of N=95±9.7 (Mean±SD) cells remain in the community after each round of bottlenecking of this magnitude (Fig. 3B).
2. Migrations: After the top community has been selected and replicated using a standard dilution factor (*d)* 95 out of 96 communities are subject to a round of migration (*n*_*mig*_ =10^2^). We choose this migration factor so that the number cells introduced via migration was comparable to the number of cells left over by the bottlenecking (Fig. 3C).
3. Bottlenecks + Migrations: After the top community has been selected and replicated using a standard dilution factor (*d*) 95 of 96 communities are first subject to a round of subject to an additional bottleneck (*d*_*bot*_ =10^4^). These communities and then subject to a round of migration (*n*_*mig*_ =10^2^).

To obtain a statistically sound sample size each of these approaches was repeated 100 times with 100 different random seeds (Fig 3E). Each iterative directed evolution experiment is compared to an equivalent NS line started with the same metacommunity. The metacommunity was propagated for 40 generations without selection (*S*=(1/*d)I* with *d*=10^3^×).

### Quantifying the ecological resistance of directly evolved, synthetic and no selection communities

In Figures 3B-D, We compare the function and resistance to ecological perturbations of i) a community obtained from directed evolution (DE) ii) a community constructed ‘synthetically’ and iii) the highest function community obtained from a no-selection line (*S* = *I/d* for 40 transfers). The DE community was obtained by iteratively applying directed evolution using both bottlenecks and migration (as shown in Figure 3D and described in the previous section).

To construct the ‘synthetic community’ we first take the DE community and count the number of coexisting taxa (*n*). We then combine the *n* top species in the species pool (i.e the *n* species with the highest *ϕ*_i_) into a single community. These species are introduced at equal abundance with the total abundance of the synthetic community (Σ_i_*N*_i_) being equal to the total abundance of the directly evolved community. 3 metacommunities are simulated, one consisting of 96 copies of the DE community, one made up of 96 copies of the synthetic community and one made up of 96 copies of NS community. All three metacommunities are allowed to equilibrate for 20 generations (Fig 4B). The maximum function at this point *F*_max_ is recorded. At generation 20 the 3 metacommunities are each subject to an identical round of directed evolution using migration from different regional species (*n*_*mig*_ =10^2^; see section on Directed Evolution). The metacommunity is then grown for another 20 generations without selection (*S* = *I/d*).

*F*\* denotes the function of the new stable community that forms after the perturbation. We quantify the ecological resistance (*R*) as the deviation of community function after a pulse or press perturbation, as defined in [79].

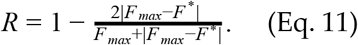

The resistance *R*ϵ[−1,1], where *R*=1 if the community function has not changed after the perturbation. Note that the resistance defined here does not reflect the sign of functional changes (e.g., increased or decreased function due to perturbation). In Figure 4C and 4D we show the *F*\* and *R* of the 95 compositional variants for each of the three communities considered.

We can calculate the overall resistance of a single community as the Mean(*R*) and Mean(*F*\*) across all of the 95 compositional variants (i.e ignoring the unperturbed community). To obtain a statistically sound sample size we repeat the entire procedure 100 times (i.e 100 different starting DE communities and 100 corresponding Synthetic communities and NS communities (Fig 4E-4F). In Figure S9 we repeat this whole procedure using different types of perturbations other than migration (specifically bottleneck, resource shift, and species knock out). For these simulations we use *d*_*bot*_=10^4^ and δ = 1.

## Code Availability

All simulations were conducted in python using *Ecoprospector* (https://github.com/Chang-Yu-Chang/Ecoprospector). All the data analysis was conducted in R. The complete code used for this paper can be found on the GitHub repository (https://github.com/Chang-Yu-Chang/community-selection).

## Acknowledgments

The authors wish to thank Rob Phillips and Hernan Garcia for inviting most of us, either as instructors or as students, to participate in the Physical Biology of the Cell summer course a the Marine Biology Laboratory in Woods Hole, MA, where this project was started and the first version of the *Ecoprospector* package was coded. We also wish to thank Brian Von Herzen for his input and discussion in the early phases of this project. This work was supported by the National Institutes of Health through grant 1R35 GM133467-01, and by a Packard Foundation Fellowship to AS. C-YC was supported by a graduate fellowship Government Scholarship to Study Abroad by the Government of Taiwan. MR-G was supported by a Gaylord Donnelley postdoctoral fellowship through the Yale Institute for Biospheric Studies.

