## Supplementary Materials for "Top-down engineering of complex communities by directed evolution"

This PDF includes:

- Supplementary Text
- Supplementary Figures 1-11
- Supplementary Tables 1-5

### Supplementary Text

#### Effect of infant population size on Migrant-pool and Propagule Strategies

The migrant-pool and propagule strategies were inspired by earlier group selection experiments, which were carried out with small populations (e.g. N=16 individuals at the start of each batch incubation) of sexually reproducing and genetically diverse animals (20, 33–36). The combination of small population sizes and sexual recombination ensures a sufficiently large between-population variation, on which group-level selection can act. By contrast, microbial populations are largely clonal and they are much larger (e.g. $10^7$-$10^9$ cells/mL is commonplace). For instance, in the original experiment by Swenson et al, the inoculum consisted of 0.06-6 g of soil, which should generally contain no fewer than ~$10^6$ and up to ~$10^{10}$ bacteria (aside from other microorganisms) (56). The simulations reported in Fig. 1E-F were inoculated with $10^6$ cells, and this number is representative of typical population sizes at the beginning of every batch. Given the large population size and clonal reproduction in our communities, we reasoned that the migrant-pool and propagule strategies may be limited in their ability to generate between-community variation in function, and thus will fail to improve community phenotypes even when communities are stable.

Consistent with this idea, we found that pooling the top-performing communities of a stable metacommunity generally increases the mean $F$, but decreases $F_{max}$ (Fig. S5A-B). This is partly due to lower-contributing (low $\phi_i$) species coming from low function communities outcompeting the high-contributing (high $\phi_i$) species in the top community (Fig. S6). Importantly, pooling the top-performing communities dramatically suppresses between-community variation in the offspring generation (Fig. S5B Inset), rapidly exhausting the ability of artificial selection to act (28). This caveat has been raised before when the migrant pool was applied to animal populations (20), but it is exacerbated here likely due to the large inocula that are common in microbial community selection experiments, and which we have replicated in our simulations. As for propagule propagation (Fig. S5C-D), when applied to stable communities the community-level heritability (which quantifies the degree to which the function of offspring communities resembles their parents' (19)) is very high, approaching $h^2$~1.0 in most simulations (Fig. S7). This high heritability explains the strong response of the mean $F$ to propagule selection (Fig. S5D-inset). Unfortunately, given the large population sizes and the fact that our species reproduce asexually, high heritability implies that the best community after selection is very similar to the best community in the parent population, both compositionally and functionally (30). Propagule strategies can thus be efficient at preserving community function, but when they are combined with a high infant population size do not introduce enough variation to improve it much beyond that point (Fig S8A-D). Consistent with this idea we find that both of these methods can work when the infant population is much smaller, i.e when they are combined with a harsh bottleneck (Fig S10).

### Supplementary Figures

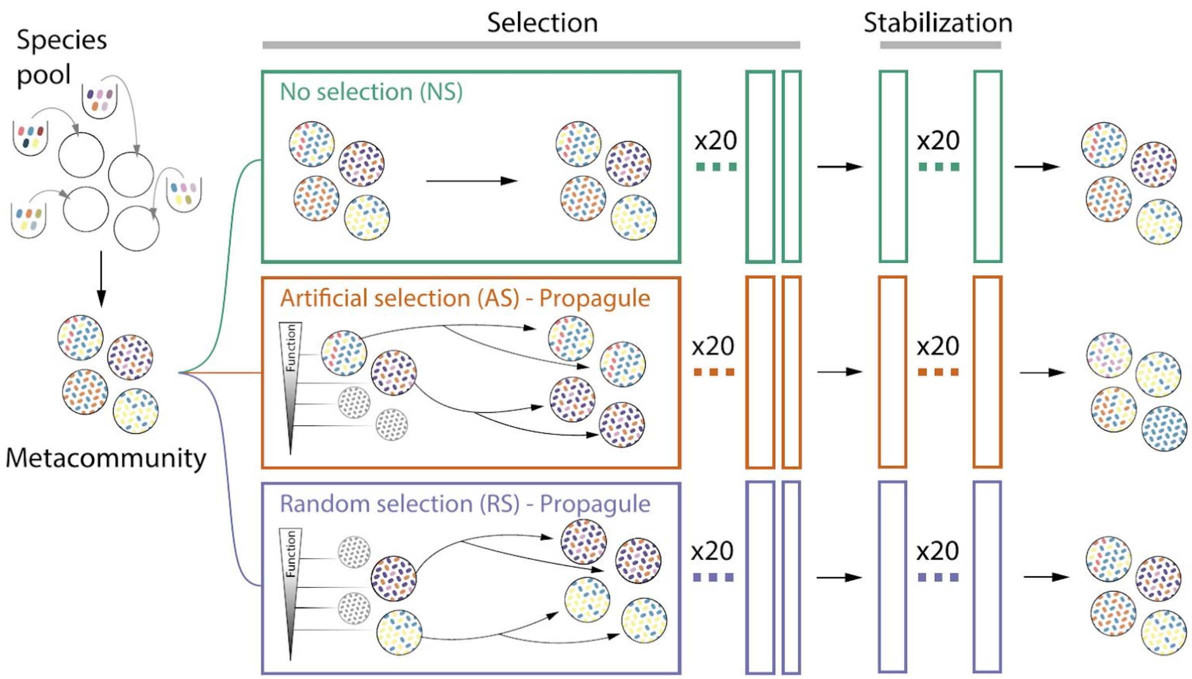


**Supplementary Figure 1. Propagule strategy of microbial community selection.** Depiction of a commonly used experimental artificial selection (AS) protocol (the propagule Method (20, 33)), alongside its two controls, no selection (NS), and random selection (RS).

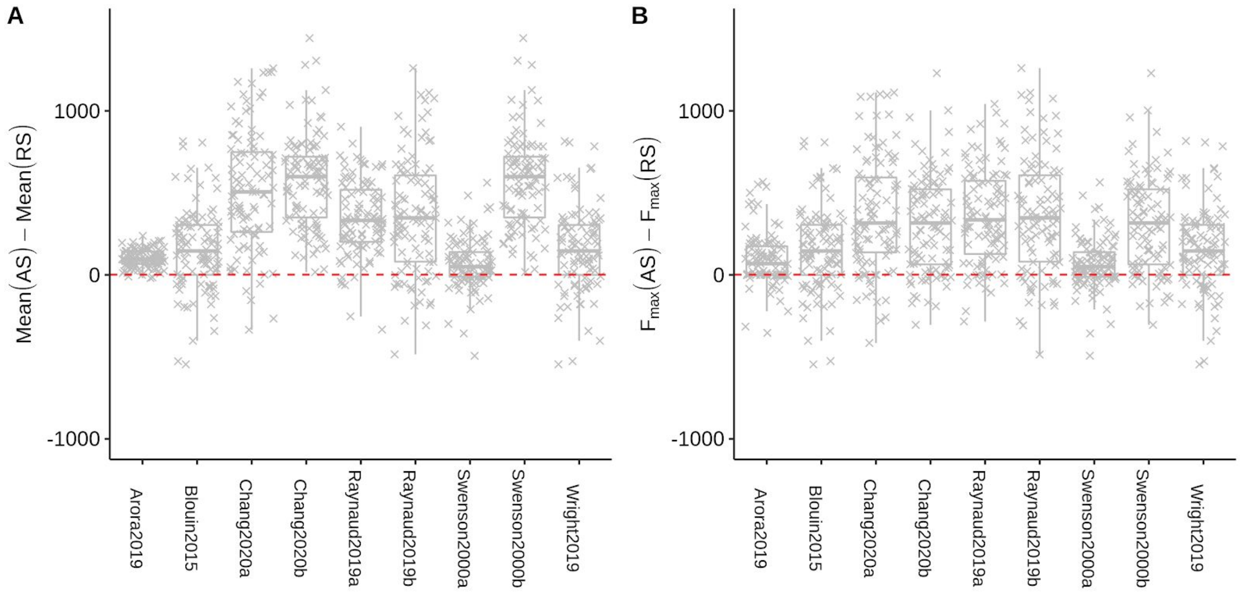


**Supplementary Figure 2. Mean and maximum function of artificial selection (AS) line relative to the random selection line (RS).** Difference in (**A**) mean function and (**B**) $F_{max}$ between the AS and RS lines. Only experimental protocols that have described RS are shown (Table S1). All differences are statistically significant (Welch's t-test, $P < 0.01$, $N = 100$).

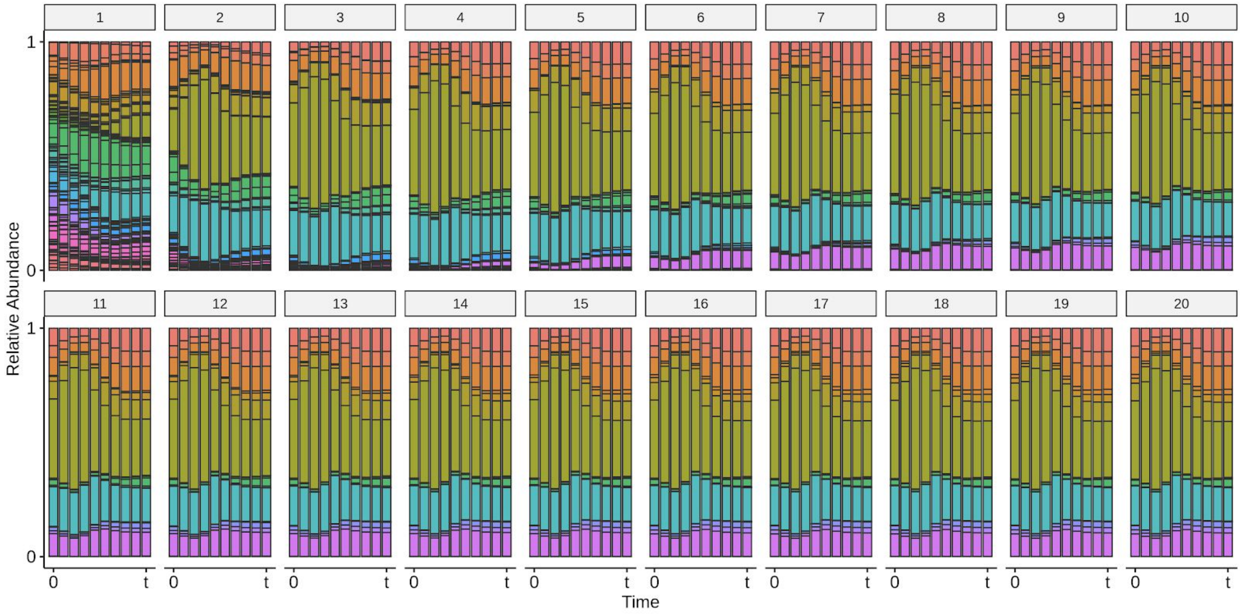


**Supplementary Figure 3. Internal dynamics of a community within a generation.** To illustrate the concept of generational stability we plot the within-batch community dynamics of a single community in the no-selection line over 20 generations. Each vertical bar represents one of 10 time points within a growth cycle, and colors represent taxa. The initial inoculum has 228 taxa, most of which rapidly go extinct in the first five growth cycles. Despite the temporal dynamics changes within each growth cycle, after ~10 generations community composition converges to a dynamic equilibrium reflected in a repeatable ecological succession in consecutive generations.

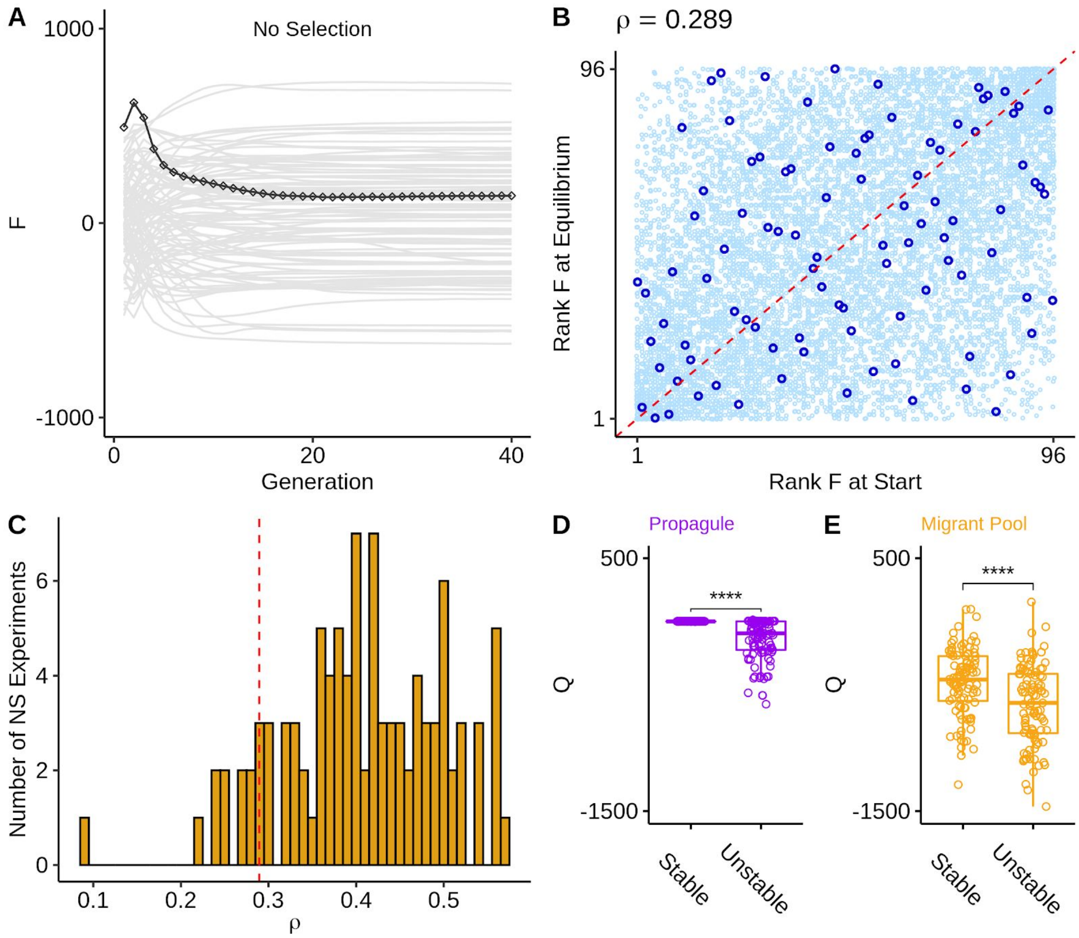


**Supplementary Figure 4. Selecting for transient communities is inefficient. (A)** Functions of 96 communities grown for 40 generations without selection (NS). Highlighted line shows the community with the highest function at the end of the first generation **(B)** Community rank function at the start of an experiment is a poor predictor for the rank function at equilibrium. Dark blue points represent 96 communities shown in panel **A**, and the light blue points denote the communities in the other 99 replicate NS lines. We calculated Spearman's $\varrho$ between Rank F at Start and Rank F at equilibrium for this dark blue points **(C)** Distribution of Spearman's $\varrho$ for all 100 NS lines. Dotted red line corresponds to Spearman's $\varrho$ for the 96 communities shown in panel **A**. **(D-E)** To confirm that selecting for transient communities reduces the effectiveness of both migrant pool and propagule selection methods, we compare an experiment where we apply 20 generations of artificial community level selection to newly inoculated metacommunity with an experiment where we apply 1 single round of artificial community level selection on a metacommunity that has already been stabilized for 19 generations. After this the metacommunity is grown for another 20 generations without selection so that the communities reach equilibrium. In panel (D) we use the propagule method and select 25% of communities after each generation. In panel (E) we use the migrant pool method and also select 25% of communities after each generation. Each of these experiments is repeated 100 times and their effectiveness compared to the NS control is quantified by $Q=F_{max}[AS]-F_{max}[NS]$. For both the propagule (D) and migrant pool (E) methods we find that a single round of selection on a set of stable communities does better than 20 rounds of selection starting on unstable communities. Brackets represent paired t-tests (N=100 for each test). ****:$p<0.0001$.

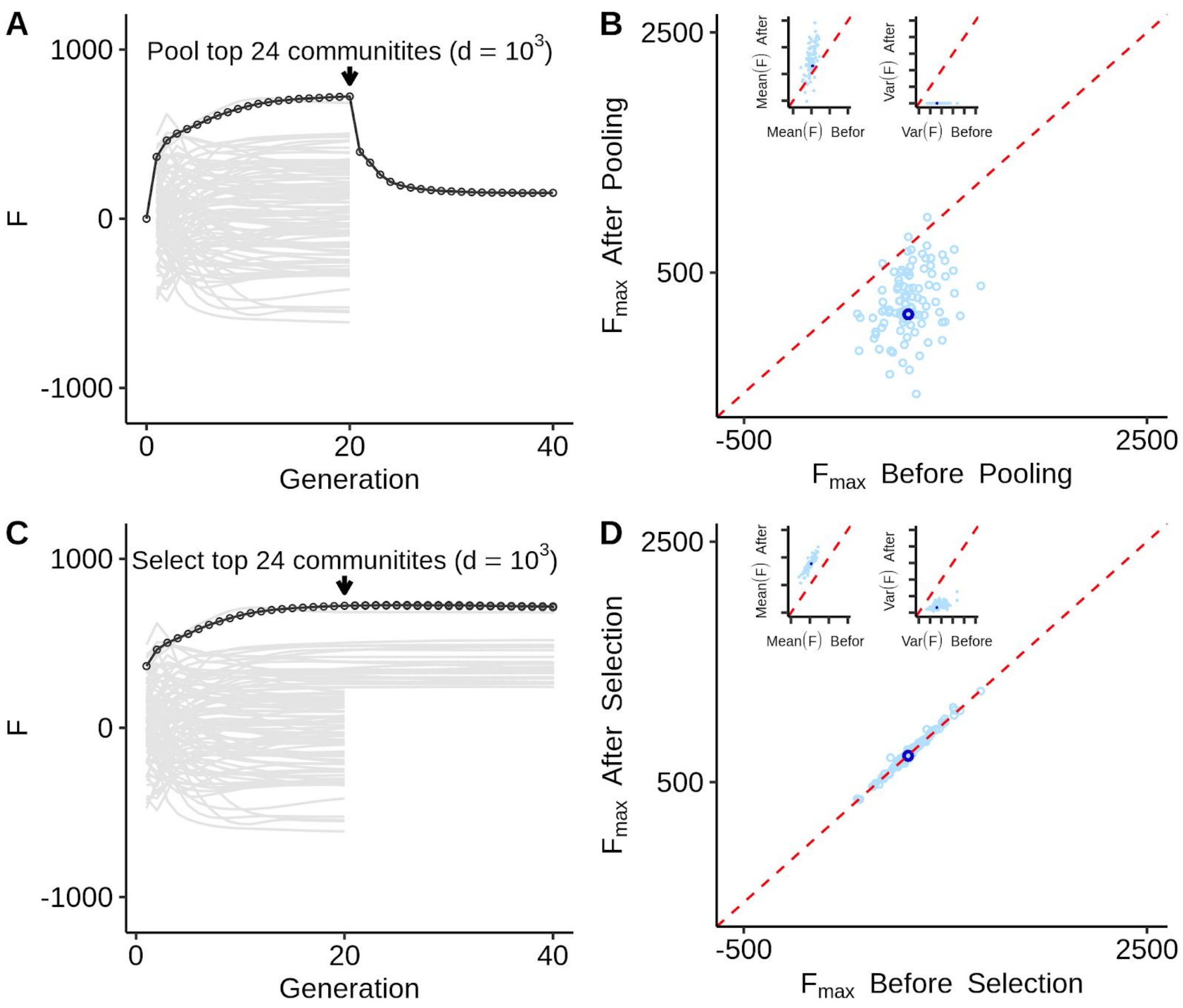


**Supplementary Figure 5 Migrant pool and propagule methods fail due to high infant community population size.** **(A)** The community with the highest function at transfer 20 (solid dark line, circular points) may drop when pooled with lower functioning communities (grey lines). (**B)** Pooling the functionally distinct communities into a single inocula usually results in higher mean function (left inset) but results in lower maximum function and reduced functional variation (right inset). Dark blue point represents the one experiment shown in panel **A**. **(C)** Selecting the top 25% of communities at transfer 20 using the propagule method with a modest dilution factor preserves the function of the top community at (sold dark line, circular points). This is due to high heritability of community function when communities are at equilibrium (Fig S7). **(B)** High heritability means that the propagule strategy consistently results in higher mean function (left inset). However it also means that the propagule strategy is unable to generate new functional variation (right inset) and so we see minimal change in the maximum function before and after selection. Dark blue points represent the one experiment shown in panel **(B).**

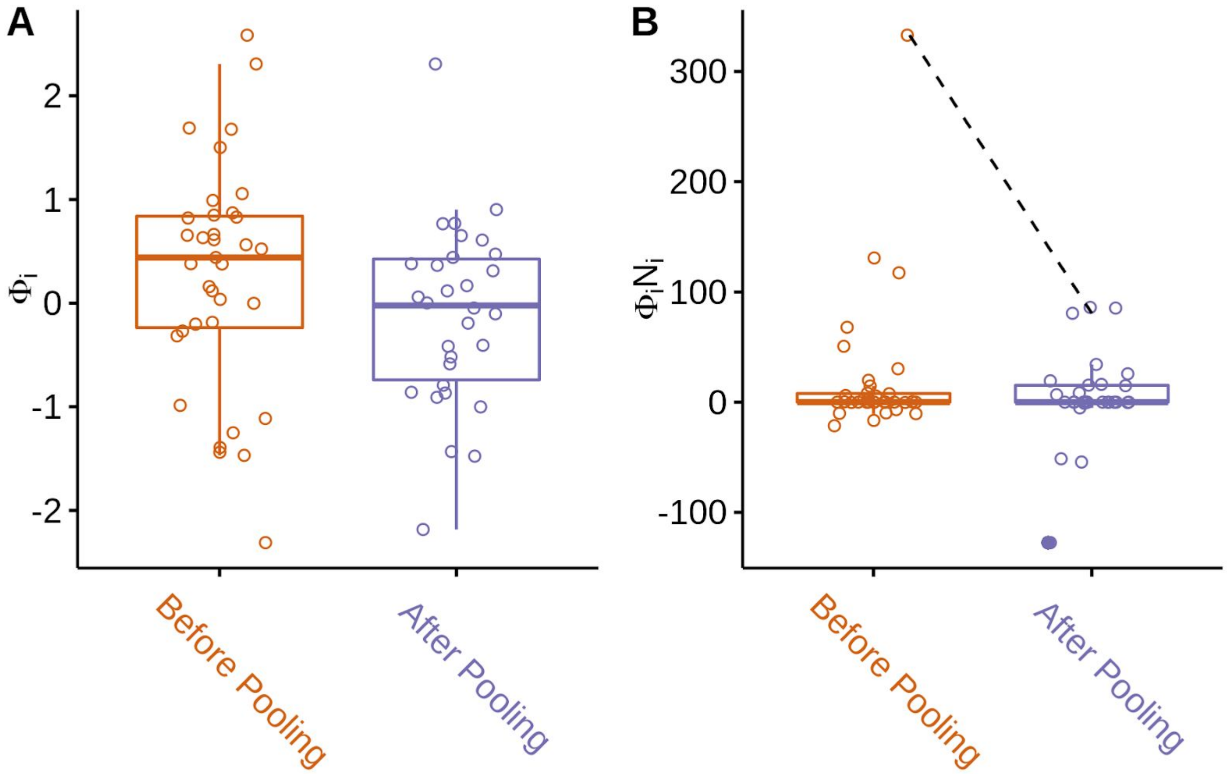


**Supplementary Figure 6. Species contribution to the function of the top community selected using migrant-pool strategy.** Each box represents the top one of 96 communities and each point represents a species in the community. Before pooling (species richness = 35) and after pooling (species richness = 30) is shown in (**A**) per-capita species contribution $_i$ and (**B**) per-species contribution (per-capita species contribution times species biomass $_iN_i$). The drop in community function ($F = \sum_i {}_iN_i$) from 723 to 152 is in part due to a substantive drop in abundance of the highest performing taxa ($N_i$ = 144 before pooling vs $N_i$ =35 after pooling). The drop in abundance is shown in panel B with the dashed line. This drop is due to competition with lower functioning new migrants introduced from other communities (such as the taxa highlighted in solid).

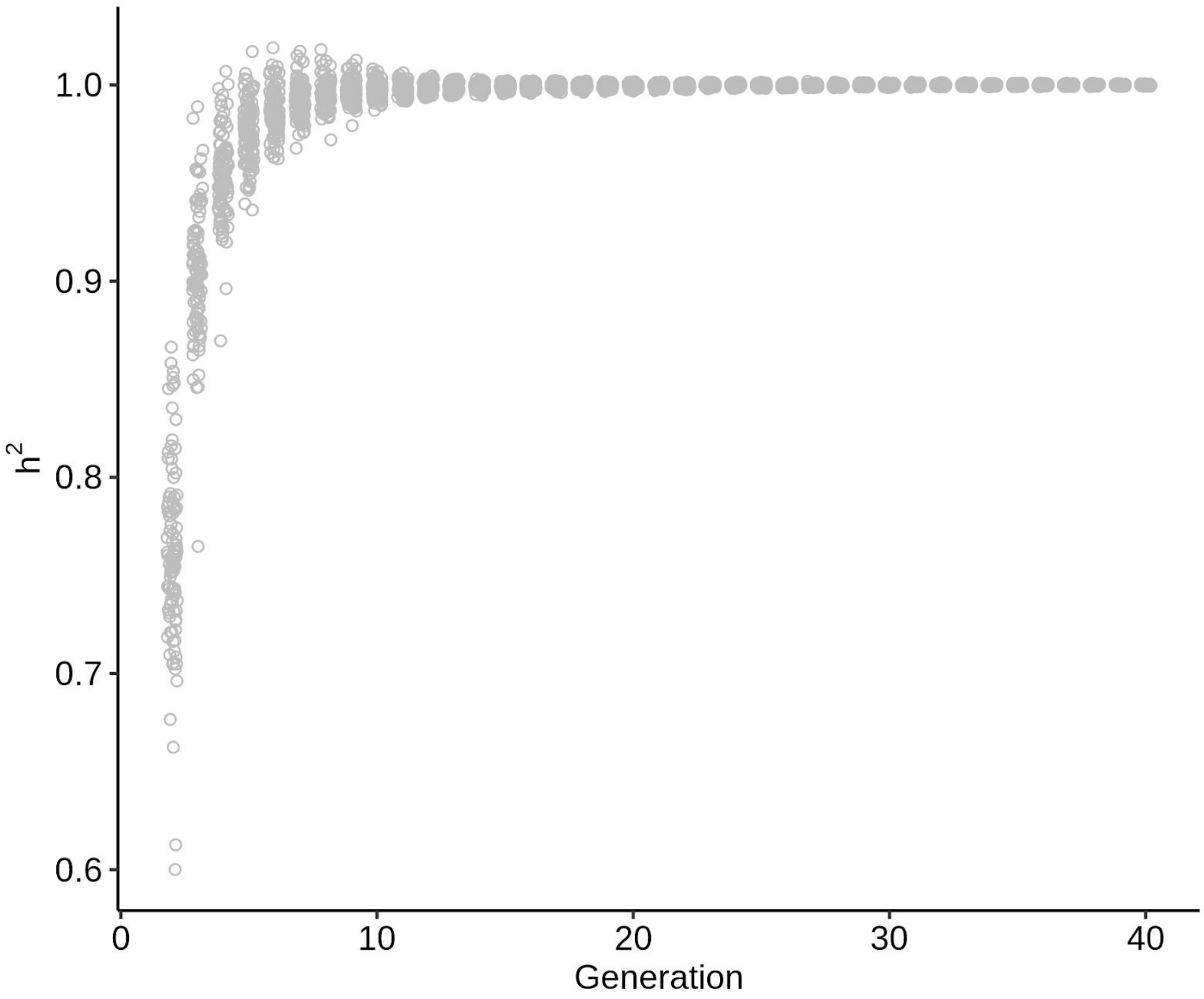


**Supplementary Figure 7. Heritability as a function of time in 100 no-selection lines.** Each point is the heritability in community function calculated using 96 parental and offspring communities. Heritability is estimated by the slope of linear regression between parental and offspring community function.

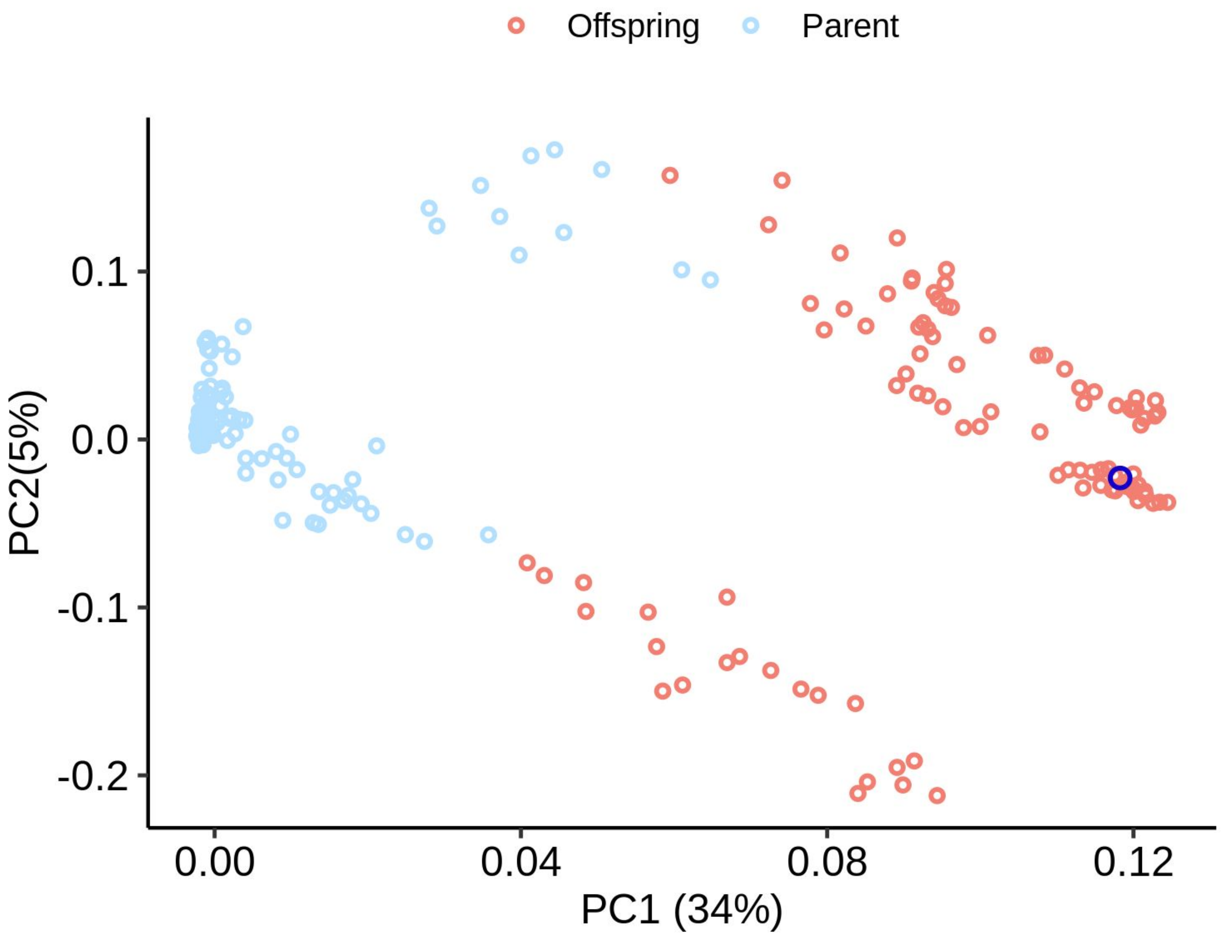


**Supplementary Figure 8 Compositional variants generated using dilution shocks.** Principal component analysis of the species relative abundance for the communities shown in Fig 2B. Light blue circles correspond to the 96 communities at the end of generation 20 (parent). Red circles correspond to the 96 communities at the generation 40 (offspring). The dark blue circle corresponds to the highest functioning community at generation 20 (i.e the one that is used to seed the offspring generation).

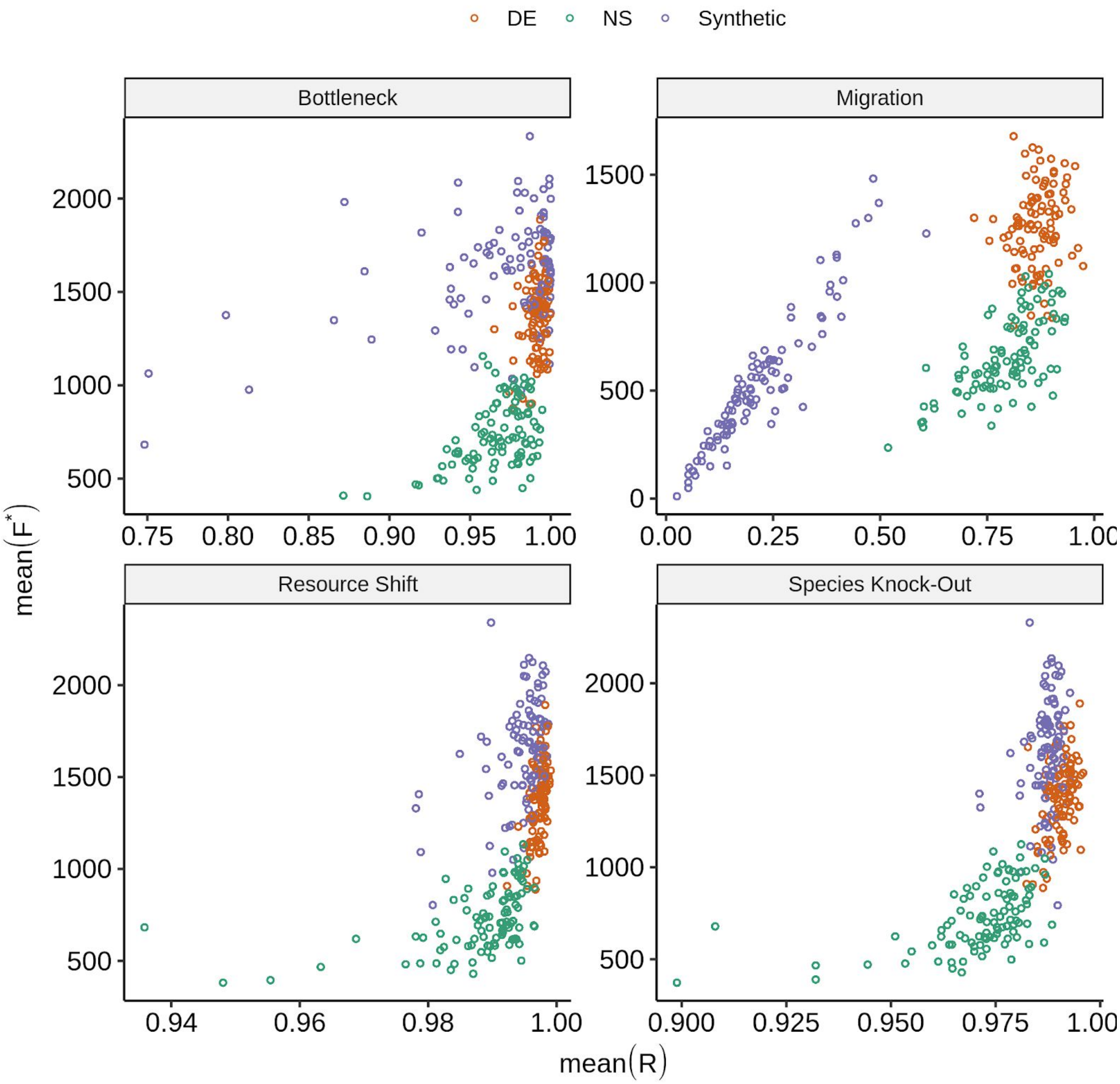


**Supplementary Figure 9. Community resistance to various types of ecological perturbation** Each subplot shows the Mean($R$) vs Mean($F^*$) for 100 independent experiments where we subjected the three types of communities described in Figure 4A to 95 replicates of single type of perturbations. The top right panel is the same as Figure 4F where the ecological perturbation examined was migration ($n_{mig}$=$10^2$). We repeat this experiment for bottlenecks ($d_{bot}$=$10^4$) (top left) , resource shifts ($\delta$ = 1) (bottom left), or species knock-outs (bottom right).

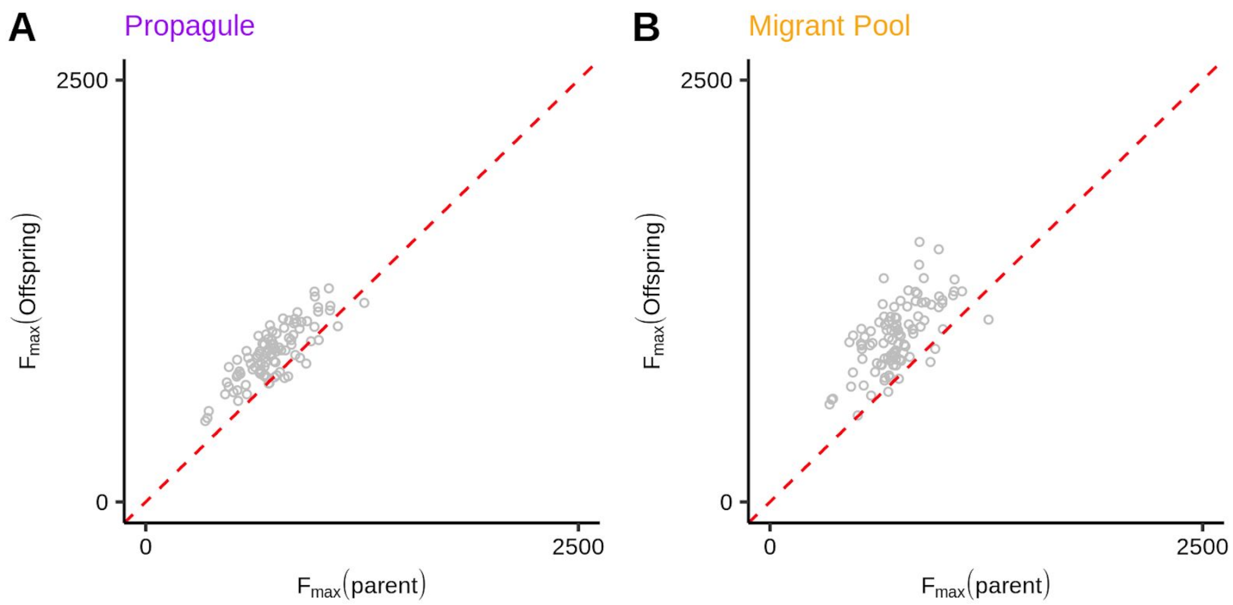


**Supplementary Figure 10. Propagule and migrant-pool approaches can improve maximum community function when a harsh bottleneck is applied.** In an experiment, a metacommunity of 96 communities is passaged for 20 generations without selection. At generation 20, 24 communities are selected and passaged according to either a propagule selection strategies (A) or a migrant pool selection strategy (B). Immediately after selection a harsh dilution shock is applied to all communities. For (A) we apply a $10^5$ bottleneck whereas for B we apply a $2x10^6$ bottleneck which means we end up with an average of ~10 cells in panel A and ~12 cells in panel B.. The communities are subject to another 20 serial transfers.

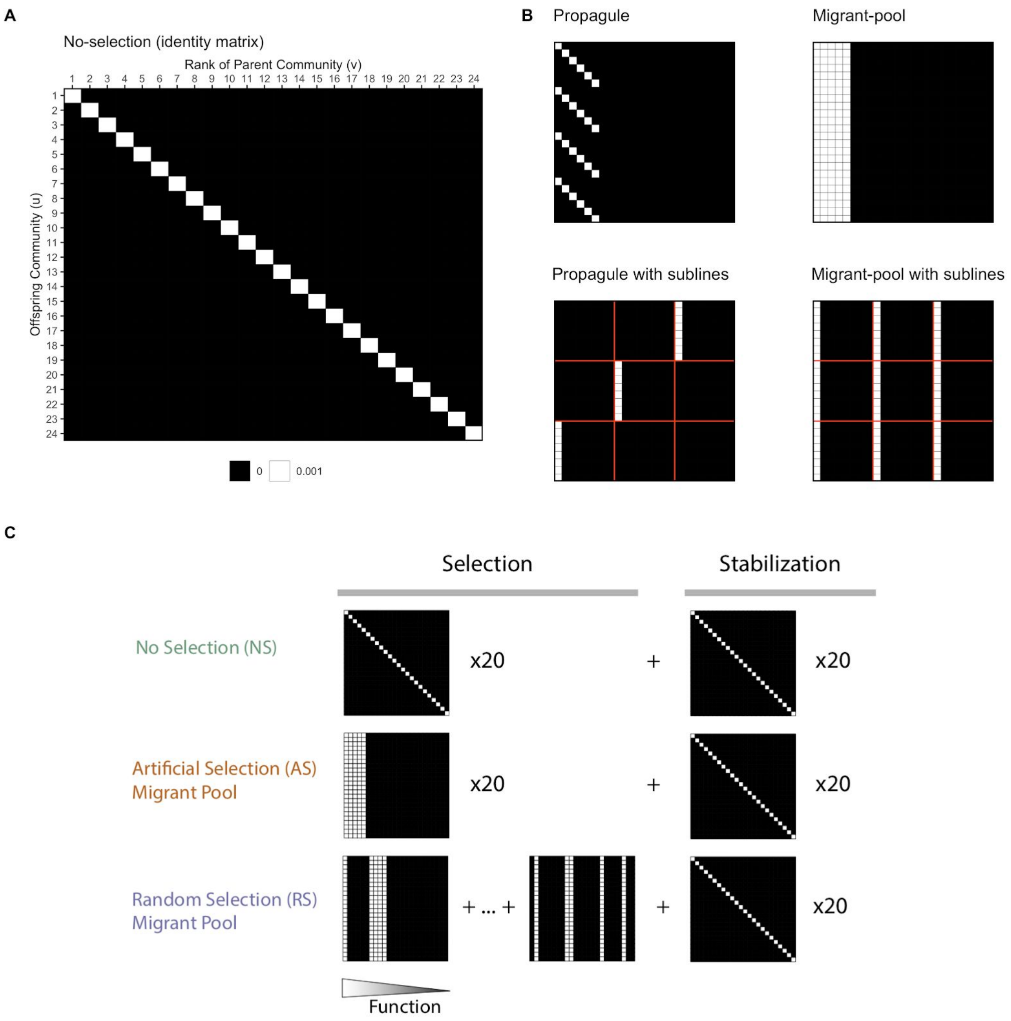


**Supplementary Figure 11. Selection matrices**. (**A**) The selection method used at the end of each generation can be encoded as selection matrix $S$ whose element $S_{uv}$ represents the fraction parent community of rank $u$ that is transferred to offspring $v$. The value of $S_{uv}$ is set to the dilution factor $d = 10^3$ that is used in batch culture. The parental communities are ordered according to their functions such that the top performing community will locate at the leftmost column in S whereas the community with lowest function will be on the rightmost. An identity matrix represents a one-to-one transfer of 24 communities without selection. (**B**) The two widely used community selection strategies in the microbiome selection studies: propagule and migrant-pool approaches can be represented as selection matrices. The fraction of parental communities being selected to seed the offspring community is $\rho = 0.25$ (6 of 24 communities) in both cases. The two matrices on the bottom show examples of protocols with three sublines [Raynaud2019], where red lines separates the 24 communities into three sublines of eight communities. (**C**). Sequence of selection matrices used for the three lines shown in Figure 1B (adapted from [Swenson2000])

**Supplementary Table 1. Experimental protocols on artificial community selection**. Seven protocols fall into the category of migrant-pool strategy in which a selected set of communities is pooled into a single inoculum to seed the next generation of communities. The other five protocols use the propagule strategy where a selected set of communities are propagated asexually to generate the offspring. Three protocols [Raynaud2019a, Raynaud2019b, and Arora2019] have sublines, which are represented as blocks separated by the red lines. The selection schemes of 12 experimental protocols are converted into selection matrices, which was used to simulate the protocols *in silico* using *Ecoprospector*.

| Strategy | Protocol | Community source | Targeted function | Random selection control | Number of generations | Number of selection lines | Number of community per lines | Number of communities selected each generation | Percentage of selected communites | Selection matrix[a] | Random selection matrix[a] |
|---|---|---|---|---|---|---|---|---|---|---|---|
| migrant pool | Swenson2000a | plant-associated soil | soil microbiome, plant host biomass | yes | 16 | 2 | 15 | 3 | 0.20 | 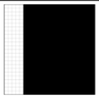 | 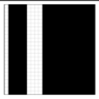 |
| | Blouin2015 | water treatment plant | lowest CO2 emmision | yes | 20 | 6 | 30 | 3 | 0.10 | 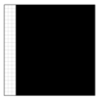 | 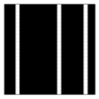 |
| | Panke-Buisse2015 | plant-associated soil microbiome | early or late flowering time in the plant host | not available | 10 | 1 | 14 | 4 | 0.28 | 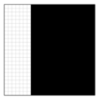 | |
| | Jochum2019 | soil microbiome associated to dought-persistent grass | delayed onset of drought stress symptom in wheat seedlings establishment | not available | 6 | 1 | 50 | 5 | 0.10 | 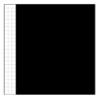 | |
| | Raynaud2019b | topsoil | biomass estimated by OD | yes | 14 | 1 | 30 | 3 | 0.10 | 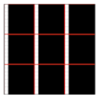 | 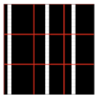 |
| | Mueller2019 | rhizospheres | biomass of plant host with salt-stress tolerance | not available | 9 | 5 | 8 | 2 | 0.25 | 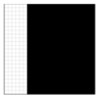 | |
| | Wright2019 | bulk marine debris | highest chitnase activity | yes | 7 | 1 | 30 | 3 | 0.10 | 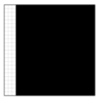 | 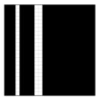 |
| propagule | Swenson2000b | aquatic microbiome from a pond | the highest or the lowest water pH | yes | 40 | 1 | 24 | 6 | 0.25 | 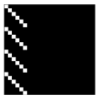 | 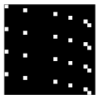 |
| | Arora2019 | fruitfly gut | shortest host eclosion time | yes | 4 | 10 | 3 | 1 | 0.33 | 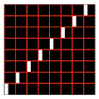 | 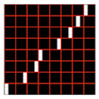 |
| | Raynaud2019a | topsoil | biomass estimated by OD | yes | 14 | 3 | 10 | 1 | 0.10 | 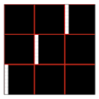 | 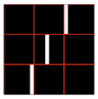 |
| | Chang2020a | synthetic communities with four known strains | amylolytic acitivities | yes | 17 | 1 | 24 | 4 | 0.16 | 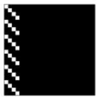 | 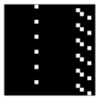 |
| | Chang2020b | soil and leaves | cross-feeding potential | yes | 7 | 1 | 92 | 23 | 0.25 | 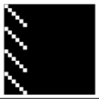 | 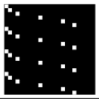 |

[a]For illustration convenience, the seletion matrices shown here are designed for 24 communities rather than 96 that are used otherwise in the main text.
[a]In our simulation, a new random selection matrix for a protocol is drawn every time it needs to transfer from parents to offsprings, so they differ from generation to generation.

**Supplementary Table 2. Parameters for Microbial Consumer-Resource Model.** Adapted from Marsland2020 Table 1.

| Parameter | Description and units | Value |
|---|---|---|
| $N_i$ | population density of species i (individuals/volume) | [a] |
| $R_\alpha$ | Concentration of resource α (mass/volume) | [a] |
| $C_{i\alpha}$ | Uptake rate per unit concentration of resource α by species i (volume/time) | [b] |
| $D_{\alpha\beta}$ | Fraction of byproducts from resource β converted to α (unitless) | [bc] |
| $g_i$ | Conversion factor from energy uptake to growth rate (1/energy) | 1 |
| $w_\alpha$ | Energy content of resource α (energy/mass) | 1 |
| $l_\alpha$ | Leakage fraction for resource α (unitless) | 0 |
| $m_i$ | Minimal energy uptake for maintenance of species i (energy/time) | 0 |
| n | Hill coefficient for functional response (unitless) | 2 |
| $\sigma_{max}$ | Maximum input flux (mass/time) | 1 |

[a]Values change with consumer-resouce dynamics.

[b]Values are assigned randomly to each species during simulation setup.

[c]The values in $D_{\alpha\beta}$ do not matter because $l_\alpha$is 0

**Supplementary Table 3. Parameters for MiCRM.** Most parameters are adapted from Marsland2020 Table 2 except for a, scale, $n_{inoc}$, and $\alpha$.

| Parameter | Description and units | Value |
|---|---|---|
| M | Number of resources | 90 |
| T | Number of resource classes | 1 |
| H | Number of microbial species in global pool | 2100 |
| $R_{tot}$ | Total resource abundance | 1000 |
| $S_f$ | Number of specialist families | 1 |
| $u_c$ | Mean sum over a row of the preference matrix $c_{i\alpha}$ | 10 |
| $\sigma_c$ | Standard deviation of sum over a row of the preference matrix $c_{i\alpha}$ | 3 |
| $c_0$ | Low consumption level for Binary $c_{i\alpha}$ | 0 |
| $c_1$ | High consumption level for Binary $c_{i\alpha}$ | 1 |
| q | Fraction of consumption capacity allocated to preferred resource class | 0[a] |
| s | Sparsity of metabolic matrix | 0.2[b] |
| $f_w$ | Fraction of secreted byproducts allocated to waste resource class | 0.45[a] |
| $f_s$ | Fraction of secreted byproducts allocated to the same resource class | 0.45[a] |
| a | Exponent parameter in power-law distribution that determines the species abundance in regional pool | 0.01 |
| ψ | Number of cells when $N_i$ = 1 | 1e+06 |
| $n_{inoc}$ | Number of cells in the initial inoculum | 1e+06 |
| α | Relative functional contribution of species interaction to the additive case | 1 |

[a]These values do not matter if $S_f$ is 1

[b]This value does not matter if $l_\alpha$ is 0

**Supplementary Table 4. Protocol-specific parameters.** These parameters are used in the protocol used to systematically evaluate the selection matrices from empirical studies (Figure 1E-F; Table S1).

| Parameter | Description and units | Value |
|---|---|---|
| d | Dilution factor in the batch culture | 0.001 |
| t | Incubation time | 1 |
| $n_{wells}$ | Number of wells; number of metacommunities | 96 |
| $T_{tot}$ | Number of total transfers (generations) | 40 |
| $T_{selc}$ | Number of selection transfers (generations) | 20 |

**Supplementary Table 5. Parameters for directed selection in Figure 2D-2I.**

| Parameter | Description and units | Value |
|---|---|---|
| $\theta$ | The percentile determining the high-performing species in the species pool used to knock in | 0.95 |
| $d_{bot}$ | Bottleneck size | 1e+05 |
| $n_{mig}$ | Number of cells in the migrant community | 1e+06 |
| $f_{coa}$ | Mixing ratio of coalescence; biomass of immigrant community relative to that of a perturbed community copy | 0.5 |
| $\delta$ | Tunes the magnitude of resource perturbation. The fraction from depleting a resource and move the same amount to another | 1 |
